# Homology-aware average nucleotide identity estimation at scale with TurboANI

**DOI:** 10.64898/2026.11.05.764740

**Authors:** Jianshu Zhao, Daniel McDonald, Lucas Patel, Claire Wang, Antonio Gonzalez, Rob Knight

## Abstract

Here we introduce TurboANI, a fast homology-aware average nucleotide identity (ANI) estimation algorithm that implements the *seed-chain-quasi-alignment* scheme using SIMD seeding, compact indexing, lightweight chaining, and cache-friendly bitset sliding minwise hashing, achieving up to ∼40,000-fold speedup over the alignment-based ANI algorithm with near-perfect agreement. TurboANI is also >40-fold faster than the state-of-the-art FastANI and ten-fold faster than skani while maintaining better accuracy. In the sensitive mode, it has a ten-fold higher true-ANI recovery rate than skani.

## Main text

Large collections of bacterial, archaeal and viral genomes (e.g., metagenome-assembled genomes, or MAGs) representing diverse ecological and evolutionary histories from a wide range of environments are now publicly available ^1–7^. This deluge of genomic data provides an opportunity to robustly investigate important questions in microbial ecology and evolution, including medical applications such as the human microbiome and bacterial pathogenesis. At the same time, it underscores the need to advance existing methods for analyzing these large-scale genomic datasets. One of the primary tasks among these methods is to compute whole-genome similarity, or average nucleotide identity (ANI), which serves as the fundamental step for species and intraspecies unit delineation ^8–11^, new species/strain identification and homologous recombination analysis ^12, 13^, and other tasks. Classical methods for computing the similarity and homologous regions are based on sequence alignment; for example, OrthoANI ^14^ and JSpecies ^15^ are based on USEARCH ^16^ and BLAST ^17^, respectively. However, these methods cannot scale to more than a handful of genomes ^18^. FastANI, a sketching-based method, has been among the most highly cited metagenomics/genomics papers since publication (>6,200 according to Google Scholar) due to its fast algorithm design and efficient implementation, achieving up to 1,000-fold speedup over alignment-based methods ^19^. Despite its efficiency, FastANI does not scale to contemporary genome collections. This has created a computational challenge, bottlenecking many downstream applications including genome similarity-based database search (a key step for genome classification) ^20^, genome taxonomy classification ^21^, genome clustering to define biological units ^22^, strain profiling ^23^, genome assembly polishing ^24^ and many widely used metagenomic analyses through platforms such as MiGA^25^, Anvi’o ^26^ and KBase ^27^.

FastANI first splits the query genome into fixed-length fragments (default 3 kilobase pairs, or 3 kbp) and extracts random minimizers from both these query fragments and the reference genome by hashing k-mers with the 128-bit MurmurHash3, then keeping the smallest canonical hash per sliding window (here we refer to it as the minimizer window). The minimizer window length and the bottom-s sketch size are not arbitrary constants: FastANI derives them from the k-mer length, query fragment length, minimum identity threshold, and *P*-value cutoff so that the sampled minimizers are sufficient to detect mappings at the target identity while limiting random matches. For each query fragment, the L1 stage looks up its minimizers against the reference minimizer index using the lower 64 bits of the 128-bit hashes and flags reference regions with enough matching minimizer seeds as coarse candidate intervals, each roughly the length of the query fragment. The L2 stage then refines each candidate interval with a sliding-window bottom-s minwise hashing (or MinHash): the query fragment is walked across the window one position at a time, incrementally updating a running bottom-s sketch as it slides, and at each position the Jaccard similarity against the fragment’s sketch is computed, with the best-scoring position kept as that fragment’s mapping and its Jaccard index converted into a Mash-distance identity estimate ^28^. This L1/L2 process runs independently for every query fragment, and the final ANI is the average of the per-fragment identity estimates across all fragments with a mapping (the best mapping only). This process (via the Mashmap algorithm ^29^) is closely related to long-read mapping, a well-studied problem popularized by minimap2 ^30^, which follows a seed-chain-extension framework ^31^ with theoretical guarantees ^32, 33^. FastANI differs from seed-chain-extension schemes ^31^ used in general-purpose long-read alignment because it does not perform chaining or base-level alignment extension. Instead, minimizer seed hits are used to identify candidate reference intervals, although anchor chaining can naturally be introduced at this stage, as shown in minimap2 ^30^ or skani ^34^. FastANI then applies a quasi-alignment step, based on sliding-window bottom-s MinHash over minimizer hashes ^29^, for homologous region identification and identity estimation. Thus, before the final alignment or quasi-alignment step, both FastANI-like ANI estimation and minimap2-like read mapping must obtain minimizer seeds and identify candidate reference intervals for each query sequence. Only selected candidate intervals are passed to the expensive final step: either base-level extension alignment in read mapping or sliding-window MinHash quasi-alignment in FastANI ^29, 30^.

To further scale ANI calculation for modern genomic datasets, we reformulated the sketch-based ANI estimation problem as a *seed-chain-quasi-alignment* problem and developed TurboANI. TurboANI implements the *seed-chain-quasi-alignment* scheme (Fig. 1 and Methods): SIMD minimizer extraction accelerates minimizer seed generation (Fig. 1a and Methods); twisted tabulation hashing provides fast, high-quality minimizer hashes (Fig. 1a and Methods); a compact flat inverted index reduces reference-index memory use and cache misses (Methods); diagonal clustering plus lightweight chaining reduces L1 candidate intervals (Fig. 1b,c and Methods); a bitset-based dynamic order-statistic structure accelerates the L2 sliding window bottom-s MinHash (Fig. 1d and Methods); and reusable precomputed Poisson distance tables remove repeated distance and confidence-bound calculations during identity estimation (Methods). These algorithmic and systems-level improvements jointly reduce unnecessary window evaluations and improve cache locality by replacing pointer-heavy, general-purpose tree-based bookkeeping with compact array- and bitset-based data structures.

**Figure 1.**
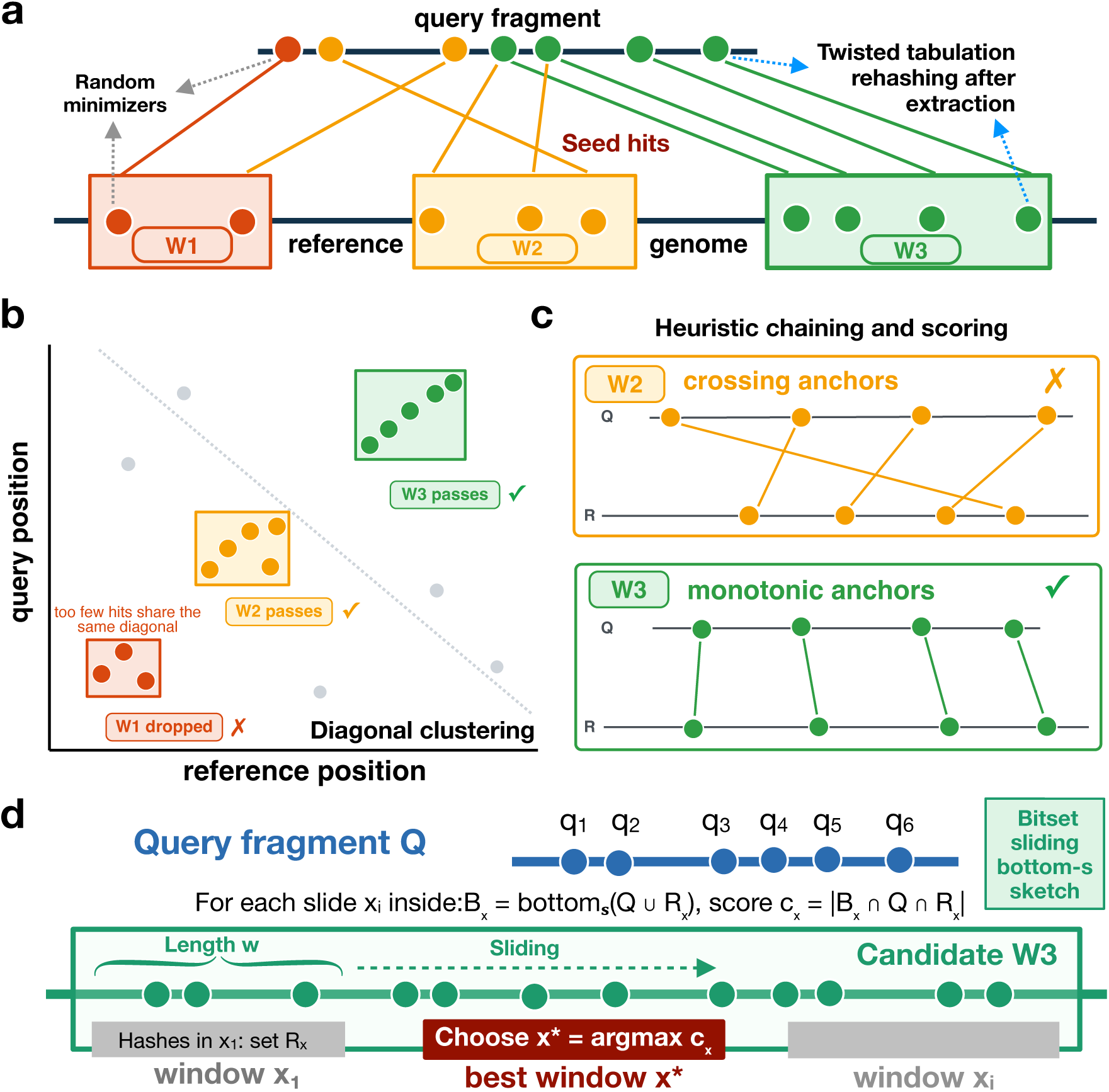
Schematic overview of the seed-chain-quasi-alignment framework in TurboANI. **a**. SIMD minimizer seeding and rehashing via twisted tabulation hashing. Each shape represents a k-mer hash while the position is the minimizer position. The highlighted intervals on the reference sequences are initial intervals identified by matched minimizers only. **b**. Diagonal clustering of query fragments and the reference genome by examining the relative positions of matched minimizer positions for each initial interval. **c**. Minimap2-style fast and heuristic chaining of candidate intervals that pass the diagonal clustering filtering in **b**. **d**. Bitset sliding window MinHash over candidate intervals that pass the chaining step in **c**. In L2, each reference window spans a fragment-length interval in minimizer-window-start coordinates, and the bottom-s score is evaluated over the union of query-fragment minimizer hashes and reference-window minimizer hashes. The best window is selected by maximizing the bottom-s objective function.

TurboANI speedup over a single thread scales nearly linearly with increasing thread count (Fig. 2a), consistent with the highly parallel structure of the algorithm. In the current implementation, query genomes are mapped in parallel against a read-only reference index, and within each query genome, query fragments are also mapped in parallel. On a collection of ∼12,000 genomes spanning 75% to 100% true ANI (Fig. S1) based on alignment-based methods, TurboANI’s fast mode is around 41 times faster than FastANI and around 10 times faster than skani-sensitive at a similar true ANI recovery rate (Fig. 2b, 2c, Fig. S2 and Fig. S3, see Methods for a detailed definition of ANI recovery rate). TurboANI’s default mode is >20 times faster than FastANI and also has several times larger true ANI recovery rate than FastANI and skani-sensitive using OrthoANIu as the ground truth (Fig. 2c, Fig. S2 and Fig. S3). When BLAST-ANI was used as the ground truth, TurboANI’s default mode had a >10-fold true ANI recovery rate than skani-sensitive and is as good as FastANI (Fig. S4). TurboANI’s sensitive mode is around 7-fold faster and also >10-fold more accurate than FastANI. The sensitive mode is also >10 times more accurate than skani-sentitive while being as fast as skani-sensitive (Fig. 2c). Mean absolute error (MAE) analysis showed that the TurboANI sensitive mode is significantly more accurate than skani-sensitive (MAE=1.32% and 2.47%, respectively, Fig. S5) based on OrthoANIu ANI truth. Mean absolute error (MAE) analysis based on BLAST-ANI truth showed that TurboANI’s default mode is in near perfect agreement with the truth and is significantly more accurate than FastANI and skani-sensitive (Fig. 2d, 2e and 2f). When compared with FastANI for each of the steps in the *seed-chaining-quasi-alignment* framework, TurboANI is around 142 times faster for the seeding step (or the minimizer extraction step), around 10 times faster for the candidate interval screening step (or the L1 step), around 48 times faster for the sliding window MinHash step (or the L2 step) and 443 times faster for the distance calculation step (Fig. 2g and 2h). To test if a different seeding strategy would affect TurboANI accuracy and speed in addition to random minimizers, we implemented open and closed syncmers ^35^. However, both open and closed syncmers are less accurate than random minimizers when compared to the ground truth (Fig. S6a) and can be 2-4 times slower than random minimizers despite SIMD support (Fig. S6b). To further understand the chaining component that is new in TurboANI, we counted the number of candidate intervals (L1) that go into the L2 quasi-alignment step with or without the diagonal clustering and chaining step using a 300-genome collection from the bacterial family *Streptomycetaceae*. Compared to FastANI, diagonal clustering itself reduced the number of L1 intervals by 20.5% while both diagonal clustering and minimap2-style lightweight chaining (the default option) reduced the number of L1 intervals by 35.8% (Fig. 2i). If a more expensive and accurate co-linear chaining algorithm, optimized ChainX, is combined with diagonal clustering, the number of L1 intervals can be reduced by 42.9% (Fig. 2i). However, the overall program runtime and peak RAM increased by 18.7% and 15.1%, respectively (Table S1) and the ANI estimation accuracy was almost identical to minimap2-style lightweight chaining (Fig. S7). Therefore, diagonal clustering together with minimap2-style chaining was chosen as the default L1 candidate interval screening algorithm. This default chaining step reduces the number of L2 sliding windows by 132 billion compared to the original FastANI on the 300-genome testing dataset, serving as the key algorithmic improvement that makes L2 sliding-window MinHash faster. We then also added paired minimizer filtering, inspired by strobemers/randstrobes ^36^ (two or more short k-mers (“strobes”) sampled from nearby windows along a sequence and concatenated together, rather than taken as one contiguous substring) after the default L1 candidate interval screening, further reducing the number of L1 candidate intervals by 41%. However, it is considerably less accurate despite being 1.3 times faster (Table S2). We then examined the number of candidate intervals for a chosen query fragment throughout the default L1 candidate interval screening process. The results showed that only three candidate intervals were retained from 22 intervals from initial seed hits after diagonal clustering and one candidate interval was retained after minimap2-style chaining (Fig. S8). Therefore, there is only one candidate interval left for the L2 step for this query fragment (Fig. S8). Our new bitset sliding MinHash algorithm is around 4.6 times faster than the default ordered map-based implementation when the same number of sliding windows were used (Fig. 2j).

**Figure 2.**
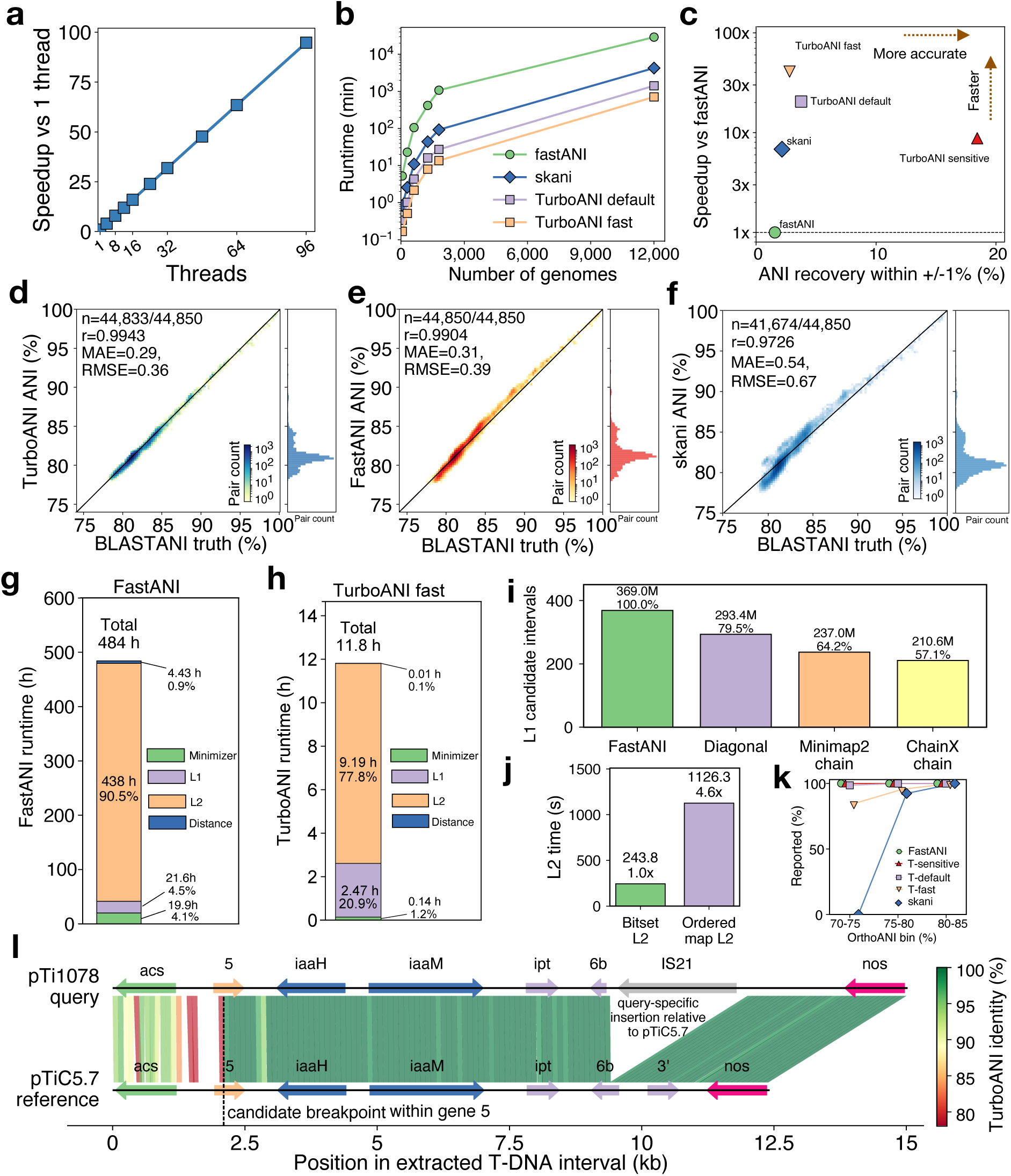
Speed and accuracy benchmark of TurboANI against FastANI and skani. **a**. Speedup of TurboANI over a single thread as the number of threads increases (running time for 96 threads is extrapolated). **b**. Runtime of TurboANI default mode, fast mode, skani-sensitive and FastANI against the number of genomes. **c**. TurboANI (three modes) and skani-sensitive speedup over FastANI versus true ANI recovery rate based on OrthoANIu (Methods). **d**. Correlation, MAE and RMSE analysis between TurboANI default mode and BLAST-ANI. **e**. Correlation, MAE and RMSE analysis between FastANI and BLAST-ANI. **f**. Correlation, MAE and RMSE analysis between skani sensitive mode and BLAST-ANI. **g**. Runtime of each individual seed-chain-quasi-alignment step in TurboANI using the 12,000-genome collection. **h**. Runtime of each individual seed-quasi-alignment step in FastANI using the same 12,000-genome collection. **i**. Number of candidate intervals obtained by each L1 algorithm for the 300-genome collection. **j**. Runtime comparison for the bitset sliding MinHash and ordered-map sliding MinHash. **k**. Number of reported ANI values by TurboANI, skani and FastANI for individual ANI bins from 75% to 85%. **l**. Homology map from TurboANI of the query Ti plasmid pTi1078 against the reference plasmid pTiC5.7.

We then benchmarked TurboANI against skani, the fastest sketch-based ANI estimation algorithm. For the 300-genome collection, skani-sensitive reported 91% of the pairs for ANI between 75% and 80% and 0% of the pairs between ANI 70% and 75% when compared to ground truth (Fig. 2k). However, TurboANI’s fast mode reported 100% and 83% for the two ANI ranges and reported 100% for all ANI ranges in the default and sensitive mode (Fig. 2k). Analysis based on mean absolute error (MAE) within each OrthoANIu bin showed that skani-sensitive was less accurate than TurboANI default and sensitive modes across nearly all ANI ranges, and was better only than TurboANI fast mode in the 70-75% ANI range (Fig. S9a). Analysis based on residual standard deviation showed that skani-sensitive had >2 times greater pair-to-pair variability than TurboANI modes, with TurboANI sensitive mode showing the lowest variability, followed by TurboANI default (Fig. S9b). For residual variability, TurboANI fast mode was more stable than FastANI above 90% ANI, whereas FastANI was more stable than TurboANI fast mode below 85% ANI (Fig. S9b). Analysis based on mean absolute error (MAE) within each BLAST-ANI bin showed consistent results with the OrthoANIu bin (Fig. S10a and S10b). For the default skani and ANI > 85%, our experiments showed that it is as fast as TurboANI fast mode but is also less accurate than TurboANI fast mode (Table S5). Together, these results indicate that skani had considerably less stable estimation errors than TurboANI, even when their average errors were similar for some large ANI pairs (e.g., > 90% ANI). Next, we benchmarked genome database search and taxonomy classification accuracy using the same 300-genome collection and OrthoANIu as the truth. TurboANI achieved substantially higher nearest-neighbor recall than skani-sensitive, improving mean recall by approximately 10 percentage points for both top-5 and top-10 retrieval (Fig. S11). BLAST-ANI based nearest-neighbor recall analysis showed consistent results with OrthoANIu-based analysis (Fig. S12). Because nearest-neighbor search is a rank-sensitive task, this improvement indicates that TurboANI more accurately preserves local genomic similarity neighborhoods relative to OrthoANIu and BLAST-ANI truth (note that for nearest neighbor search tasks, 10% percentage improvement is substantial ^20, 37^). The lower skani recall suggests that its ANI estimates introduce enough pair-to-pair variability to perturb the ordering of the closest database genomes, consistent with the above MAE and residual standard deviation analysis. We then explored the use of rapid ANI calculation for clustering genomes into biological units (e.g., species or strains) and compared the greedy clustering accuracy of TurboANI against FastANI and skani-sensitive using orthoANIu as the ground truth. For the same 300-genome collection, TurboANI default and sensitive modes were consistent with OrthoANIu clustering across ANI thresholds (Fig. S13a). The greatest advantage was at the 85% ANI and lower threshold, where TurboANI sensitive and default recovered 83 and 77 clusters compared with 95 OrthoANIu clusters, whereas FastANI and skani-sensitive recovered only 64 and 65 clusters. At the species-level threshold of 95% ANI, TurboANI default and sensitive recovered 223 clusters compared with 224 OrthoANIu clusters, while FastANI recovered 217 and skani-sensitive recovered 229 (5 wrong clusters); at the strain-level threshold of 99.5% ANI, all TurboANI modes and FastANI exactly recovered the 291 OrthoANIu clusters, whereas skani-sensitive recovered 292. Cluster representative assignments showed the same pattern (Fig. S13b): at 85% ANI, TurboANI sensitive and default assigned 268/300 and 242/300 genomes to the same longest-genome representative as OrthoANIu, compared with 212/300 for FastANI and 183/300 for skani-sensitive, while at 95% and 99.5% ANI clustering thresholds the representative assignments were nearly identical across methods, with TurboANI matching 298-300/300 genomes. The same clustering accuracy analysis based on BLAST-ANI showed consistent results with OrthoANIu (Fig. S14a and S14b). We then benchmarked TurboANI against skani and FastANI using a collection of 280 archaeal genomes from the family *Methanobacteriaceae*. We observed similar results regarding ANI estimation accuracy compared to the 300-genome collection based on OrthoANIu (Fig. S15, S16 and S17). We also compared TurboANI with skani-sentitive and FastANI for virus ANI calculation using either OrthoANIu or BLAST-ANI as truth, similar ANI estimation accuracies were observed (Fig. S18).

The L2 sliding-window quasi-alignment step in TurboANI follows the FastANI/Mashmap framework and preserves the genomic coordinates of the best-scoring reference window for each mapped query fragment. As an illustrative case, we analyzed the Ti plasmid pTi1078 from *Arminella tumorigenes* strain 1078 to show how TurboANI fragment-level homology maps can nominate local regions of shared ancestry, mosaic structure, or candidate recombination for downstream evolutionary analysis. First, we queried this plasmid against PlasmidDB ^38^ using GSearch ^20^. For the top 10 closest plasmids based on overall Mash distance, we then compared the query plasmid with the top 10 reference plasmids to find the plasmid that has the highest ANI to the query plasmid using TurboANI. A homologous T-DNA interval from pTiC5.7 of *Martinezella rhizogenes* strain C5.7 was found for the Ti plasmid pTi1078. The query was the 15.0-kb pTi1078 T-DNA acs-to-nos interval (position 144409-159433), and the reference was the 12.4-kb pTiC5.7 acs-to-nos interval (position 178404-190821). The query interval contains the genes acs, gene 5, iaaH, iaaM, ipt, 6b, an IS21 element, and nos, whereas the reference contains acs, gene 5, iaaH, iaaM, ipt, 6b, gene 3’, and nos (Fig. 2l and Extended Data. S1). The TurboANI ribbon map showed extensive collinear homology between the two T-DNA regions, but also revealed a mosaic structure. The left part of the interval, spanning acs and the 5’ part of gene 5, showed lower and more variable identity, with TurboANI fragment identities as low as ∼79% (Fig. 2l). In contrast, the region downstream of the breakpoint within gene 5 was highly conserved, with most fragments from gene 5 through iaaH, iaaM, ipt, 6b, and nos showing very high identity, often near 100% (Fig. 2l). TurboANI also identified a query-specific interval corresponding to the IS21 insertion in pTi1078, which lacked a corresponding homologous segment in the pTiC5.7 reference (Fig. 2l). The results were further validated by independent BLAST-based homology analysis (Fig. S19a and S19b). The conclusion was unchanged despite some small identity differences for the genes around the breakpoint gene 5.

The sampling density for forward sampling schemes (e.g., minimizers are a type of forward sampling) determines TurboANI memory consumption, similar to the seed-chain-extension algorithm in long-read mapping. For our k-mer length and window range (k=16, w up to 24, down to 16 as the minimum identity decreases), double decycling ^39^ achieves the lowest sampling density and the open-closed minimizer ^40^ is a close second, typically only ∼1-4% higher in density than double decycling, while both outperform the random minimizer ^40^. However, double decycling is more expensive to compute because it requires an O(k)-like decycling-set test per k-mer, whereas the open-closed minimizer can be implemented with rolling hashes, syncmer classification, and sliding-window minima; therefore, the open-closed minimizer is computationally close to the random minimizer and is amenable to SIMD acceleration. An efficient SIMD-enabled open-closed minimizer implementation would require a dedicated SIMD kernel comparable in complexity to existing SimdMinimizers. Whether these lower-density schemes preserve sufficient sampling quality for robust candidate interval detection and downstream ANI estimation remains an open question. In addition to random minimizers, syncmer-based schemes—including open and closed—can improve sensitivity to more divergent sequences (e.g., at 75% sequence identity) by selecting more conserved, evenly spaced seeds ^35, 41^, consistent with theoretical analyses of local k-mer selection ^42^; however, our experiments showed that open and closed syncmers did not improve ANI estimation accuracy and are significantly slower than random minimizers (all were SIMD supported). Parameterized syncmers has been reported to be 50–100% slower than the default random minimizer scheme in minimap2 despite the fact that more reads can be mapped ^41^. More importantly, they lack efficient SIMD-amenable implementations that support efficient super-k-mer window (the longest contiguous substring of a sequence, consisting of several overlapping k-mers, all of which share the same minimizer) extraction after seeding used by TurboANI for rehashing. The minimizer Jaccard estimator used in TurboANI (and in FastANI) is theoretically biased as s k-mers (s=1 in the minimizer Jaccard estimator) must be selected within each local minimizer window to guarantee unbiasedness ^43, 44^, where s is the bottom-s sketch size derived beforehand (s>>1). The theoretical unbiased estimator is therefore called minmer Jaccard estimator. However, our experiments showed that it is significantly less accurate than FastANI and TurboANI when compared to the OrthoANIu truth (Table S3 and Fig. S20). Moreover, this approach is ∼2.0 times slower and consumes 2.4 times more RAM (Table S3), because initial candidate-interval construction, L1 screening, and L2 quasi-alignment all become more expensive. Furthermore, this minmer Jaccard sampling scheme is difficult to implement with an efficient SIMD kernel for fast seed/minmer extraction, especially for large-scale ANI comparisons.

Reducing the number of candidate intervals that go into the final sliding bottom-s MinHash step is the key speedup of TurboANI, similar to the seed-chain-extension approach in long-read mapping, where seeding and chaining play a similar role. However, chaining itself is compute-bound (e.g., dynamic programming). We also added an optimal co-linear chaining in TurboANI and it reduced the number of windows further without significant overall speedup. Even more efficient anchor chaining algorithm such as llchain ^45^ was proposed recently. However, it is only faster for > 10,000 anchors while the number of TurboANI anchors is approximately 300 (considering the 3kbp query fragment length and the sampling density of random minimizers). A fine-grained parallel chaining algorithm was recently proposed for minimap2, and it worked well for read mapping because a single long query-reference alignment can otherwise leave threads idle ^46^. However, our all-versus-all workflow fragment mapping is already saturated at the query/pair level with Rayon parallel iterators. We also explored an idea inspired by strobemers/randstrobes and linked minimizer ^36, 47^, which requires two minimizer hits with approximately consistent query/reference spacing. Because this additional screen reduced candidate windows at the cost of lower ANI recovery accuracy, especially for weak homologous fragments in the low- and middle-ANI range, it was not used in the default TurboANI workflow because it is less accurate.

TurboANI rehashes the selected minimizers after SimdMinimizers with twisted tabulation hashing before L1 candidate interval screening and L2 sliding-window bottom-s MinHash, rather than reusing the hash values used for minimizer selection. This separates two statistical roles: one hash function is used to choose minimizer positions for speed via SIMD, while an independent twisted tabulation hashing defines the bottom-s ordering used for identity estimation. This is closer to the assumptions of bottom-s MinHash, where the hash ordering is applied to a fixed set. The bottom-s MinHash sampling analysis in Thorup (2013) ^48^ shows that bottom-s error bounds can inherit concentration guarantees from the underlying hash-threshold sampling process; therefore, if the hash function gives stronger concentration for threshold counts, bottom-s sampling inherits stronger tail bounds. Twisted tabulation is attractive in this setting because it remains very fast but provides exponential decay in the deviation parameter *r*, down to a probability floor determined by the tabulation-table space, whereas 2-independent hashing gives only Chebyshev-style *O*(*r*^−2^), tails and *d*-wise independence gives polynomial *O*(*r*^−*d*^) tails ^48^. This stronger tail behavior is especially relevant for TurboANI because L2 evaluates many overlapping reference intervals and selects the maximum-scoring window (Methods), so rare upward hash fluctuations can otherwise be amplified into incorrect homologous-window choices (e.g., MurmurHash3 in FastANI does not have the guarantee). Our additional analysis showed that TurboANI—without diagonal clustering and minimap2-style chaining, but with SimdMinimizers and twisted tabulation rehashing—is still more accurate than the original FastANI for mid-range (85%–95%) ANIs (Table S4; Fig. S21). Therefore, the improved middle-ANI accuracy compared to FastANI is consistent with both fewer candidate intervals from the new L1 candidate screening entering L2 and a cleaner, independent bottom-s hash ordering with stronger sampling concentration from twisted tabulation hashing.

TurboANI utilizes CPU-friendly data structures including compact inverted index for initial window selection, stamped scratch arrays for allocation-light diagonal clustering, coordinate-compressed bitset sliding window MinHash and shared precomputed distance tables for fast identity calculation (Methods). Further speedups are therefore more likely to come from reducing the number of candidate intervals or reformulating the quasi-alignment problem than from additional local data structure tuning. However, our experiments with stricter L1 window filtering suggest that aggressive candidate reduction creates a speed-accuracy tradeoff, especially in the low-and middle-ANI range where weak but real homologous fragments are important. Mapping low-identity fragments fast with high accuracy, for example from 70% to 85% identity, is exceptionally challenging; this is consistent with minimap2, which maps reads with high accuracy mainly above approximately 85% identity ^30^, further supported by theoretical analysis ^32, 33^. This suggests that TurboANI is approaching an information-limited regime of the *seed-chain-quasi-alignment* framework: below this identity range, homologous fragments produce fewer reliable shared seeds, while unrelated or repetitive regions can generate competing candidate intervals. In this regime, stricter filtering removes noise but also removes true weak signal, so further speedups from candidate pruning alone are unlikely to be free. For the bitset L2 sliding MinHash, this implementation is highly cache-friendly and instruction-efficient, relying mainly on dense arrays, bit operations, set-bit search, and popcount. To our knowledge, existing ANI tools and commonly used bottom-s MinHash implementations do not use this bitset-based dynamic order-statistic formulation for sliding-window MinHash, including the most recent Mashmap 3 ^44^.

Under our benchmark conditions for both smaller ANI ranges (e.g., < 85% ANI) and larger ANI ranges (e.g. >90% ANI), skani-sensitive is less accurate than TurboANI and FastANI in terms of MAE and also showed less stable estimation errors, with larger residual variance not captured by MAE alone. It is worth noting that skani relies on supervised ANI corrections against MUMmer-based ANI. A well-known problem for MUMmer-based ANI is that it is significantly less accurate than USEARCH-based ANI and BLAST-based ANI for low ANI values, e.g., ANI 75% to 85% ^18^. We note that skani is based on sparse chaining of FracMinHash k-mers, without quasi-alignment or base-level extension alignment ^34^. This design can recover enough anchors for accurate genome-wide ANI estimation and can internally approximate homologous regions when they are long and sufficiently similar. However, because the anchors are sparsely sampled and the chain intervals are collapsed into genome-level ANI and aligned-fraction summaries, skani is not designed to provide the exact query-fragment-to-reference-window coordinates reported by FastANI and TurboANI (e.g., comparable with BLAST-based homology analysis). In contrast, although TurboANI also does not perform base-level extension alignment, its sliding-window MinHash quasi-alignment retains the best-scoring reference coordinate and local identity estimate for each mapped query fragment, providing fragment-level homology maps in addition to genome-wide ANI. This coordinate-resolved output is useful when downstream analyses require explicit homologous intervals, such as locating candidate mosaic segments, large insertions or deletions, or putative recombination breakpoints. Similar short-fragment FastANI synteny maps have been used for comparative-genomic visualization and biological interpretation; for example, Kuzmanović et al. used them to identify a putative recombination breakpoint in mosaic DNA regions ^49^. Thus, TurboANI provides a convenient way to nominate candidate local evolutionary events at much larger scales and with substantially higher throughput, while retaining accuracy comparable to BLAST-based approaches.

In conclusion, TurboANI’s contribution is less about ANI estimation per se than about showing that sketch-based genomic comparisons benefit from the same seed-chain-extend discipline that has driven long-read mapping. By recasting ANI estimation as *seed-chain-quasi-alignment* and co-designing each stage around both the statistical structure of the problem and modern CPU hardware, we close much of the remaining gap between algorithmic design and achievable throughput. As microbial genome collections continue to grow, we expect these principles — hardware-aware seeding, accuracy-preserving candidate pruning, and cache-friendly sliding window data structures — to generalize to related large-scale sketch-based tasks, including genome and metagenome database search, taxonomic placement and scalable detection of homologous genomic regions.

## Methods

### Random minimizer extraction and rehashing

The minimizer extraction step in FastANI relies on a 128-bit hash function for better randomness. However, we noticed that in forward sampling schemes ^50^, like random minimizers, samples within a small window, from which the universe of possible k-mers is always small. Therefore, a 32-bit hash function is as good as a 128-bit one in practice ^51^. More importantly, recursive string hashing ^52, 53^, as popularized by ntHash ^54^, is significantly more efficient for long DNA sequences. SimdMinimizers ^51^ hashes k-mers with a 32-bit ntHash rolling hash and identifies per-window minima using an amortized O(1) two-stacks sliding-window minimum algorithm, followed by deduplication of adjacent repeated minimizer positions. To exploit SIMD, the sequence is split into overlapping chunks that are processed as independent SIMD lanes using AVX2 or NEON, allowing several minimizer streams to be computed in parallel. This approach was reported to be 3.4-6.8x faster than the fastest non-SIMD forward-minimizer implementation and over 15x faster for canonical minimizers. TurboANI uses SimdMinimizers for efficient random minimizer extraction (canonical), obtaining both the minimizer position and its super-k-mer window for rehashing. Stronger pseudo-randomness reduces hash collisions and is important both for maintaining a low false-positive rate among candidate intervals passed to the expensive L2 stage, and for the sliding MinHash used within L2 scoring. We therefore rehash the minimizer sequence extracted from each super-k-mer window using 64-bit twisted tabulation hashing ^55, 56^, while retaining SimdMinimizers’ original position information. This is necessary because SimdMinimizers’ default 32-bit hash has a high collision probability when applied across the full set of minimizers genome-wide for candidate interval selection, rather than just the k-mers within a single minimizer extraction window. To implement open and closed syncmer-based TurboANI, we implemented a SIMD kernel for open and closed syncmer extraction in an updated version of SimdMinimizers. For the experiment on minmer, we reimplemented the minmer extraction method in MashMap 3 ^44^ using the same logic.

### Efficient candidate interval screening in L1

Minimizer hits between the query fragments and the reference in the original FastANI were used to identify potential candidate reference intervals. FastANI reports an L1 candidate reference interval when at least three shared minimizer seed hits (default options) occur on the same reference interval within approximately one query-fragment length. FastANI uses a hash-based inverted minimizer index for reference minimizer seed lookup, implemented as a general-purpose unordered_map from minimizer hash to a dynamically allocated vector of reference positions ^19^. TurboANI applies the same idea but stores the reference minimizer inverted index as a compact flat postings structure, mapping each minimizer hash to a contiguous range/array with open-addressed lookup, avoiding the pointer chasing and per-key vector overhead of a chained HashMap-style index. After candidate reference intervals are identified, TurboANI then applies two additional filtering steps: diagonal clustering and chaining. Diagonal clustering has been used in HS-BLAST ^57^, MUMmer ^58^ and minimap ^59^. The underlying assumption is that anchors from a true collinear alignment have similar reference-start minus query-start offsets, or reference-start plus query-start offsets on the reverse strand, and therefore concentrate within a narrow diagonal band, whereas unrelated hits are dispersed across many diagonals ^58^. TurboANI follows this same principle using explicit diagonal bins: anchors from a true collinear match tend to fall into the same or neighboring diagonal bins, even with modest indel drift, whereas unrelated hits scatter across many bins. Under the default L1 settings, only diagonal bins with sufficient seed support are converted into candidate reference intervals; within each selected window, the corresponding anchors are sorted by query and reference coordinates and passed to a lightweight minimap2-style chaining. In the minimap2-style chaining step, each minimizer seed hit is represented as an anchor *i* = (*x_i_*, *y_i_*, *w_i_*), where *x_i_* and *y_i_* are the reference and query end coordinates and *w_i_* is the anchor length, then sorts anchors by *x_i_*. Its chaining dynamic program is: *f*(*i*) = *max*0*w_i_*, *max*_j<*i*_[*f*(*j*) + *α*(*j*, *i*) − *β*(*j*, *i*)]8, where *α*(*j*, *i*) = *min*{*min*(*x_i_* − *x*_j_, *y_i_* − *y*_j_), *w_i_*} rewards collinear anchor extension, and *β*(*j*, *i*) penalizes gaps or diagonal changes, with *β* = ∞, if anchors are incompatible, for example *y*_j_ ≥ *y_i_* or the gap is too large ^30^. Thus, anchors that preserve order and stay near the same diagonal accumulate high chain scores, whereas scattered or inconsistent hits are penalized. Only candidate intervals supported by a sufficiently good collinear chain are passed to L2 sliding-window bottom-s MinHash quasi-alignment. We also implemented the optimal ChainX ^60, 61^ for anchor edit distance approximation between query fragments and candidate reference intervals. For each diagonal-clustered candidate, TurboANI constructs ordered anchors from shared minimizer hits and runs optimal ChainX semiglobal chaining in both forward and reverse orientations (Supplementary Methods). The algorithm seeks a collinear anchor chain with minimum gap/diagonal edit cost, using an expanding distance bound to efficiently search the feasible chaining space (Supplementary Methods). Candidate intervals supported by a ChainX chain with sufficient unique seed support are then retained for L2 sliding-window bottom-s MinHash quasi-alignment. For the paired minimizer screening after default L1, our implementation is a lightweight strobemer/paired-anchor screen after minimap2-style chaining. This resembles the linked-seed principle of strobemers/randstrobes, where non-contiguous sequence elements are combined to improve seed specificity and indel robustness, but differs from true randstrobes because the minimizers were linked after individual seed lookup rather than being precomputed and indexed as composite hash seeds.

### Cache-friendly bitset sliding-window bottom-s MinHash in L2 quasi-alignment

The L2 stage in FastANI can be viewed as a dynamic order-statistics problem over minimizer hashes. For a fixed query fragment and a candidate reference interval or window, the algorithm slides a reference window and repeatedly updates the set of reference minimizer hashes. The L2 sliding MinHash window is a fragment-length window in minimizer-window-start coordinates: for a query fragment of length *L*, TurboANI uses N = *L* − (*w* − 1) − (*k* − 1) valid minimizer-window starts: and each reference window *R_x_* contains the selected reference minimizer hashes with start positions in [x, x+N). Thus, the L2 window spans approximately one query-fragment length and contains roughly the same number of selected minimizer hashes as the query fragment. At each window position *x*, it evaluates the bottom-s sketch induced by the union of query hashes *Q* and reference-window hashes *R_x_*, *B_x_* = *bottom_s_*(*Q* ∪ *R_x_*), and counts the shared sketch elements *c_x_* = |*B_x_* ∩ *Q* ∩ *R_x_*| (Fig. 1d). The best mapping position is then selected as *x*^∗^ = *argmax_x_ c_x_* ^19, 29^. Then, *c_x_* is converted into a Mash-distance identity estimate. FastANI’s C++ implementation represents this dynamic ordered set with a balanced ordered-map structure (e.g., a red-black tree), supporting insertions, deletions, predecessor/rank-like queries, and traversal in hash order ^19^. Our bitset sliding-window bottom-s MinHash replaces this general-purpose ordered map with a local coordinate-compressed dynamic integer set. For each L2 candidate interval, all relevant hash values are known in advance: the query sketch hashes and the reference minimizer hashes in the candidate region. We sort this local universe once and replace each 64-bit hash by a dense integer coordinate (e.g. 0 to U-1 for hash universe U). Operationally, TurboANI maintains three coordinate-indexed objects for each sliding L2 comparison: a query-presence bit vector, a reference multiplicity array for the current reference window, and a union bit vector marking coordinates present in either the query or the current reference window (Fig. S22). As the reference window slides, only the coordinates of minimizers entering or leaving the window are updated; the bottom-s sketch is obtained from the first s active bits in the union bit vector, and the L2 score is the number of those selected coordinates that are present in both query and reference (Fig. S22). The sliding window then updates compact arrays and bitsets over this coordinate space rather than maintaining a pointer-heavy ordered-map structure keyed by full hash values. This idea is motivated by dynamic integer-set and mutable bit-vector rank/select data structures, which support updates together with rank, select, and predecessor-like operations over a dense universe ^62–64^. Unlike these general-purpose structures, TurboANI exploits the fact that the complete L2 hash universe is known for each sliding window, allowing the ordered hash set to be coordinate-compressed locally and updated using compact arrays and bitsets. The resulting L2 stage is closer to a streaming, cache-resident dynamic rank/select computation, which better matches the hardware behavior of current CPUs.

### Reusable Mash distance table for L2 identity estimation

After L2 identifies the best sliding-window shared-sketch count *c_x_*∗, TurboANI converts this count to ANI using a reusable Poisson Mash-distance lookup table rather than recomputing logarithms and confidence bounds for every candidate. For each query fragment with sketch size *s* = |*Q*|, TurboANI constructs or retrieves a cached table: *T_s_*[*c*], *c* = 0, …, *s*, where each entry stores both the point identity estimate and a 90% confidence upper-bound identity used for filtering. For each possible shared count *c*, TurboANI first computes 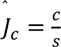, then applies the Poisson Mash transformation 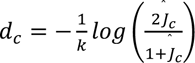 ^28^, with boundary cases *d_c_* = 1 when *Ĵ_c_* = 0 and *d_c_* = 0 when *Ĵ_c_* = 1.

The reported identity for that count is *I_c_* = 100(1 − *d_c_*). TurboANI also precomputes the relaxed identity upper bound used by FastANI-style filtering. For a distance *d_c_*, the corresponding Jaccard probability under the Poisson model is 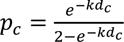. TurboANI then finds the largest integer *x* such that the upper-tail probability *P*{*X* ≥ *x*}, *X* ∼ *Binomial*(*s*, *p_c_*), is still compatible with the 90% confidence interval; equivalently, it searches until the upper-tail probability drops below 0.05, then uses *x*⁄*s* as the relaxed Jaccard value. This gives a lower bound on Mash distance, 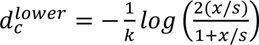, and therefore an upper-bound identity 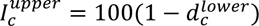. During mapping, L2 only records the best shared count *c_x_*∗. Identity estimation is then a constant-time table lookup: (*I*, *I^upper^*) = *T_s_*[*c_x_*∗]. The mapping is retained only if *I^upper^* ≥ *I_min_*. Implementation-wise, TurboANI caches one table per query sketch size *s*, so all fragments and L2 candidates with the same sketch size reuse the same precomputed Poisson distance values. This removes repeated logarithm, exponential, and binomial-tail calculations from the inner L2 loop; after the best window has been found, the distance step is reduced to indexing a small reusable table by the integer shared-sketch count.

### Selection of the minimizer window length

TurboANI adopted the same parameter-selection framework as FastANI. Given a fixed k-mer length *k*, fragment length *L*, target identity *I*, p-value cutoff *p*, alphabet size *a* = 4, and approximate reference length *R*, TurboANI follows the FastANI procedure to derive the minimizer window length. The algorithm first searches over candidate sketch sizes *s* ∈ {1,2,5,10,20, …, *L*}. For each *s*, it computes the relaxed minimum number of shared sketches *x* required at identity *I*, then estimates the probability that two unrelated sequences would share at least *x* sketches by chance. The random-match probability for a k-mer in a fragment of length *L* is 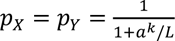, and the expected random Jaccard similarity is 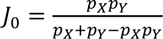. The false-positive probability for sketch size *s* is then approximated as *P_fp_*(*s*) = *R* ⋅ *P*{*X* ≥ *x*}, *X* ∼ *Binomial*(*s*, *J*_0_). TurboANI chooses the smallest candidate sketch size *s*^∗^ satisfying *P_fp_*(*s*^∗^) ≤ *p*, and converts it to a minimizer window length 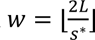, clamped to [1, *L*]. This same *w* is used when extracting minimizers from the query and reference. For the default parameters *k* = 16, *L* = 3000, *I* = 0.8, *p* = 0.001, and *R* = 5,000,000, this gives *s*^∗^ = 250 and *w* = 24. In the TurboANI fast mode, *I* is set to 0.825 while in the TurboANI sensitive mode, *I* is set to 0.775. Because TurboANI relies on SIMD, it adjusts *w* by one if necessary so that *k* + *w* − 1 is odd, which is required by the canonical minimizer tie-breaking rule in SimdMinimizer; for the default *k* = 16 and *w* = 24, no adjustment is needed.

### Parallelism via Rayon

TurboANI performs work-stealing parallel iterators at multiple levels via the Rust Rayon library (a widely used data parallelism library), including reference-genome indexing, query-genome mapping, and per-fragment L1/L2 mapping. The shared reference index is immutable, while fragment-level minimizers, candidate intervals, L2 sliding state, mappings, and counters are worker-local, avoiding lock contention in the dominant mapping stages. This design gives fine-grained load balancing for highly uneven genome-pair workloads.

### ANI truth, bacterial benchmark datasets and metrics

To obtain true ANIs for benchmarking, we use both OrthoANI (v1.2, USEARCH-based OrthoANI, or OrthoANIu) ^14^ and BLAST-ANI (or ANI_b_) ^15^ from ANI calculator (http://enve-omics.ce.gatech.edu/ani). It has been shown that OrthoANIu is essentially the same as BLAST-based ANI (ANI_b_) for ANI between 75% and 100% although more studies showed that OrthoANIu has outliers for < 85% ANI ^18^. We use two different methods as ANI truth because skani and FastANI were benchmarked differently. Skani was benchmarked against OrthoANIu while FastANI was benchmarked against BLAST-ANI. We then collected ∼12,000 genomes from the bacterial family *Streptomycetaceae* in RefSeq ^65^ and ran all-versus-all OrthoANIu or BLAST-ANI to obtain pairwise ANI truth. The ANIs from the ∼12,000 genomes are well distributed from 75% to 100% ANI. To compute the ground-truth ANI values, we distributed the pairwise comparisons across 375 compute nodes on a high-performance computing cluster, requiring approximately 6.5 million CPU-hours in total. Among these genomes, > 50% are fragmented MAGs with >200 contigs. We use default FastANI options for the benchmark. However, for skani, we cannot rely on the default settings for accuracy benchmarks because it filtered most < 85% ANI pairs without any actual sparse chaining. We therefore used a sensitivity-oriented setting with a lower identity screening threshold and slightly denser sketching so that low-ANI pairs were not discarded during the initial screening stage (id=75, c=100). This setting is more favorable to skani in the 75–85% ANI range than the faster default — a difficult range for sketch-based ANI estimation. In practice, a great majority of ANI pairs fall into that range, e.g., in a species database like GTDB ^66^. We refer to this sensitivity-oriented setting as skani-sensitive and use it throughout the manuscript unless noticed otherwise. In sketch-based ANI estimation, exact agreement with alignment-based ANI is not expected because sketching introduces sampling error and each method uses slightly different filtering and homology-detection assumptions. We therefore evaluated accuracy using an ANI recovery rate, defined as the fraction of genome pairs whose estimated ANI falls within a small tolerance of the alignment-based truth, for example 0.5%, 1%, or 2%. For an ANI bin *B*, let 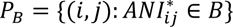 be the set of genome pairs whose alignment-based ANI falls in that bin. For tolerance *ε*, we define the ANI recovery rate as 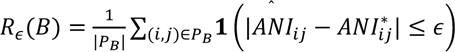, where *ANI\** is the alignment-based ANI. We report this metric across ANI bins from 75% to 100%, in addition to Pearson correlation, MAE, RMSE, and the fraction of pairs with no reported ANI. This metric is motivated by the bounded-error view used throughout sketching theory, where approximate estimators are often characterized by the probability that their error is within a specified tolerance ^67^. This bin-wise recovery analysis provides a more detailed picture of estimator behavior than a single global correlation or average error, while preventing high-ANI pairs from masking failures in the lower-ANI range. We also compute 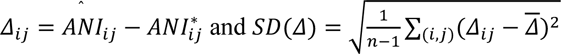 to evaluate the stability of estimation errors within each ANI bin, with larger residual SD indicating greater pair-to-pair variability after accounting for the average bias. For nearest neighbor search accuracy, recall was calculated as 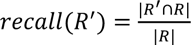, where *R*^′^ is neighbors found by TurboANI for a given genome while *R* is the true neighbors for that genome found by OrthoANIu or BLAST-ANI. For a given collection of genomes, 5% of the genomes were randomly chosen as queries and the remaining 95% as the database; this process was repeated 10 times. Average recall was reported for each repeat. All analyses were based on TurboANI v0.1.8, FastANI v1.33 and skani v0.3.2. All experiments were performed on a 64-thread Intel(R) Xeon(R) Gold 6242 CPU @ 2.80GHz with 1.5T RAM. By default, TurboANI utilizes all available threads.

### Archaea and virus genome dataset

We then randomly chose genomes from the archaeal family *Methanobacteriaceae* in RefSeq and ran all-versus-all OrthoANIu to obtain pairwise ANI truth. Complete virus genomes were retrieved from NCBI Virus database using Entrez/eUtils ^68^. To construct the broader viral benchmark, we searched for *Teseptimavirus[Organism]* AND *complete genome* and downloaded 60 complete genome FASTA records from the returned NCBI records. Viral whole-genome ANI truth was calculated using CheckV viral ANI calculator ^69^ and OrthoANIu.

### Code availability

TurboANI Rust implementation can be found in the GitHub repository: https://github.com/jianshu93/TurboANI. Scripts for reproducing the main figures/results can be found in the scripts folder of the GitHub repository. Code for additional experiments can be found on different branches of the main GitHub page. TurboANI is also available in Bioconda: https://anaconda.org/bioconda/turboani. TurboANI is also available as a Rust package here: https://crates.io/crates/turboani.

## Acknowledgements

We would like to thank Konstantinos T. Konstantinidis and Chirag Jain for their helpful comments on the manuscript. We also want to thank Ragnar Groot Koerkamp for helpful discussion on forward sampling schemes.

## Funding

This work was funded in part by the Department of Energy, USA to R.K. (DE-SC0024320). This work was also funded by Minderoo Foundation (Project title: eDNAID: Environmental DNA and AI as tools for Ocean Aid) under award number CLB-3502 and NIH under U19AG063744 (R.K.). L.P. is supported by NIH/NIGMS T32GM007198 and NIH/NIA F30AG094275.

## Author contributions

J.Z. designed the algorithm and implemented the code. J.Z. performed the software benchmarks. D.M., L.P., C.W. and A.G. helped with the software benchmarks. J.Z. wrote the manuscript with input from all authors. R.K. supervised the work.

## Competing interests

Rob Knight is a scientific advisory board member, and consultant for BiomeSense, Inc., has equity and receives income. He is a scientific advisory board member and has equity in GenCirq. He has equity in and acts as a consultant for Cybele. He is a Vice President and board member of Microbiota Vault, Inc. He is a board member of N=1 IBS advisory board and receives income. He is a Senior Visiting Fellow of HKUST Jockey Club Institute for Advanced Study. The terms of these arrangements have been reviewed and approved by the University of California, San Diego in accordance with its conflict-of-interest policies. D.M. is a consultant for and has equity in BiomeSense, Inc. The terms of these arrangements have been reviewed and approved by the University of California, San Diego, in accordance with its conflict-of-interest policies. The remaining authors declare no competing interests.

