## Supplementary Materials for "Homology-aware average nucleotide identity estimation at scale with TurboANI"

### Table of Contents

Supplementary Tables 1–5

Supplementary Figures 1–22

Supplementary Methods

### Supplementary Tables

**Table S1**. Runtime comparison between 2 different candidate interval screen algorithms for a 300-genome testing dataset. Note that for each L1 window, there are on average around 150 L2 windows per L1 window, which matched the L1 window in Figure 2f.

| L1 mode | Wall time | L2 windows | L1 CPU | L2 CPU | Peak RAM |
|---|---|---|---|---|---|
| Diagonal + minimap2-style chaining | 251.47 s | 37.08 $*10^9$ | 715.44 s | 2979.99 s | 10.98 GiB |
| Diagonal + optimal ChainX | 298.42 s | 32.87 $*10^9$ | 1857.02 s | 2598.31 s | 12.64 GiB |

**Table S2**. Paired minimizer filter as an additional L1 candidate interval screening. Default is the default diagonal clustering and minimap2-style chaining. Paired minimizer filter was added after the default screening algorithm.

| variant | wall | L1 candidates | L2 sliding windows | L2-window reduction | speedup | MAE vs fastANI |
|---|---|---|---|---|---|---|
| default | 14.08 s | 9,650,599 | 1,509,668,696 | 0% | 1.00x | 0.270 |
| Paired minimizer filter | 10.86 s | 5,690,654 | 892,915,011 | 40.9% | 1.30x | 0.418 |

**Table S3.** Minmer Jaccard estimator-based FastANI performance benchmark.

| Method | Wall time | Peak RSS | MAE | RMSE | 1% recovery | 2% recovery |
|---|---|---|---|---|---|---|
| FastANI C++ | 139.69 s | 3.32 GiB | 2.600 | 2.701 | 4.86% | 15.68% |
| Minmer FastANI | 71.28 s | 7.97 GiB | 3.915 | 4.062 | 3.88% | 5.31% |

**Table S4**. Accuracy comparions between default TurboANI, TurboANI with FastANI L1, original FastANI for ANIs between 85% and 95%.

| Method | Pearson r | MAE | RMSE | bias | +/-1% recovery |
|---|---|---|---|---|---|
| FastANI C++ | 0.9930 | 0.642 | 0.697 | +0.631 | 90.23% |
| TurboANI (FastANI L1) | 0.9944 | 0.219 | 0.280 | +0.010 | 99.67% |
| TurboANI default | 0.9947 | 0.211 | 0.269 | +0.012 | 99.73% |

**Table S5**. TurboANI fast mode versus skani default mode (fast but filters many <85% ANI pairs) for > 85% ANI only.

| Method | Truth Pairs | Matched Pairs | Matched Fraction | MAE | RMSE | Bias | Wall Time | Peak RSS |
|---|---|---|---|---|---|---|---|---|
| TurboANI-fast | 1,558 | 1,558 | 1.000000 | 0.973143 | 1.135762 | 0.941801 | 54.480 s | 4.854 GB |
| skani default | 1,558 | 1,557 | 0.999358 | 1.207078 | 1.327345 | 1.205436 | 59.950 s | 1.804 GB |

#### Supplementary Figures

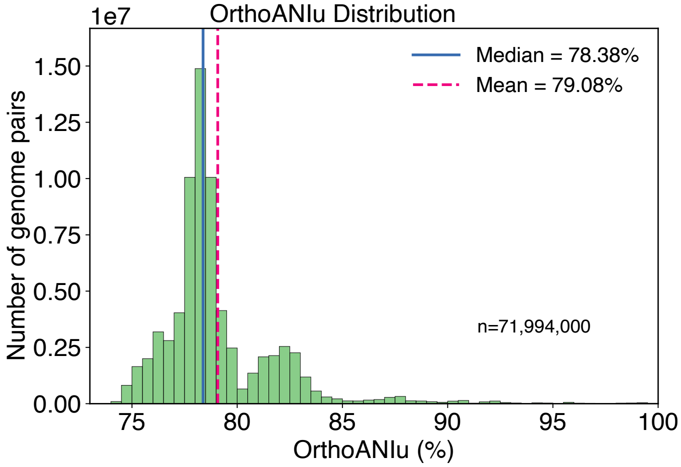


**Figure S1**. OrthoANI ANI distribution for a subsampled subset of ~12,000 genomes from bacterial family Streptomycetaceae covering ANI from 75% to 100%.

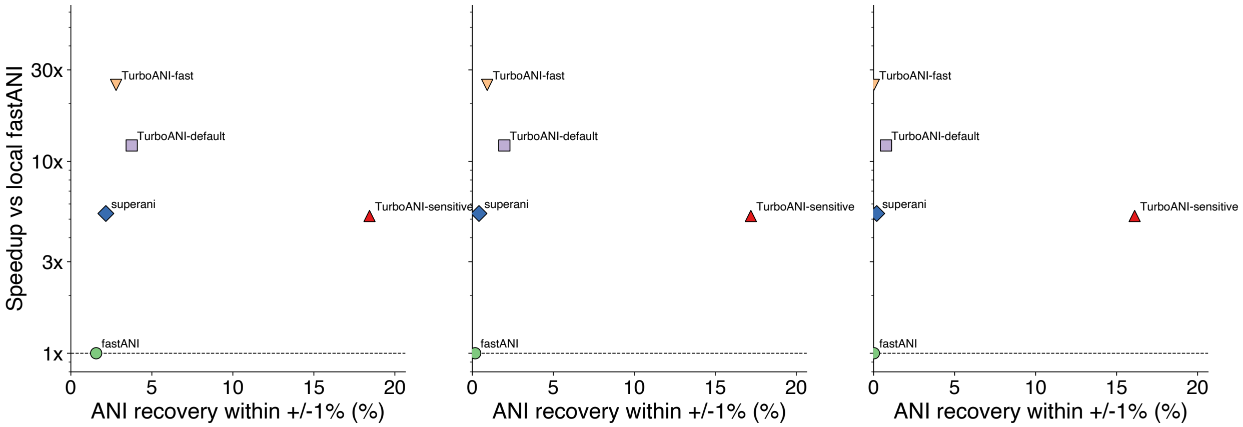


**Figure S2**. TurboANI, skani and FastANI speedup versus ANI recovery rate (within +/- 1% of the truth) against truth (OrthoANIu). (a) All ANI ranges, 75%~100%; (b) ANI range 75%~90%; (c) ANI range 75%~85%

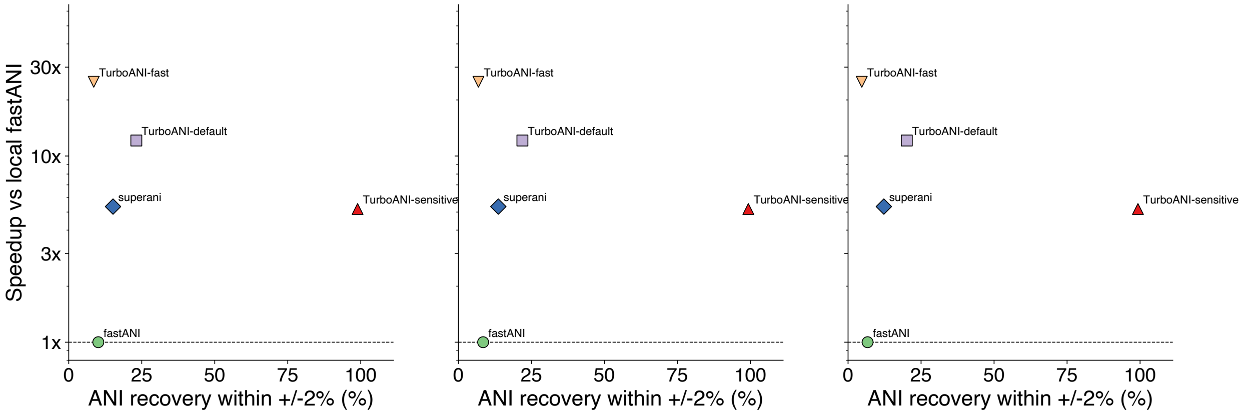


**Figure S3**. TurboANI, skani and FastANI speedup versus ANI recovery rate (within +/- 2% of the truth) against truth (OrthoANIu). (a) All ANI ranges, 75%~100%; (b) ANI range 75%~90%; (c) ANI range 75%~85%

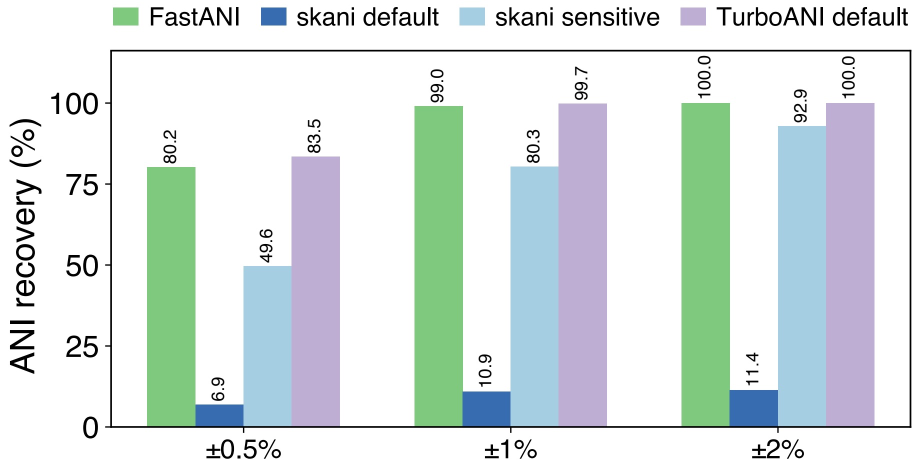


**Figure S4.** TurboANI, skani and FastANI true ANI recovery rate against BLAST-ANI truth for within +/- 0.5%, +/- 1% and +/- 2% ANI.

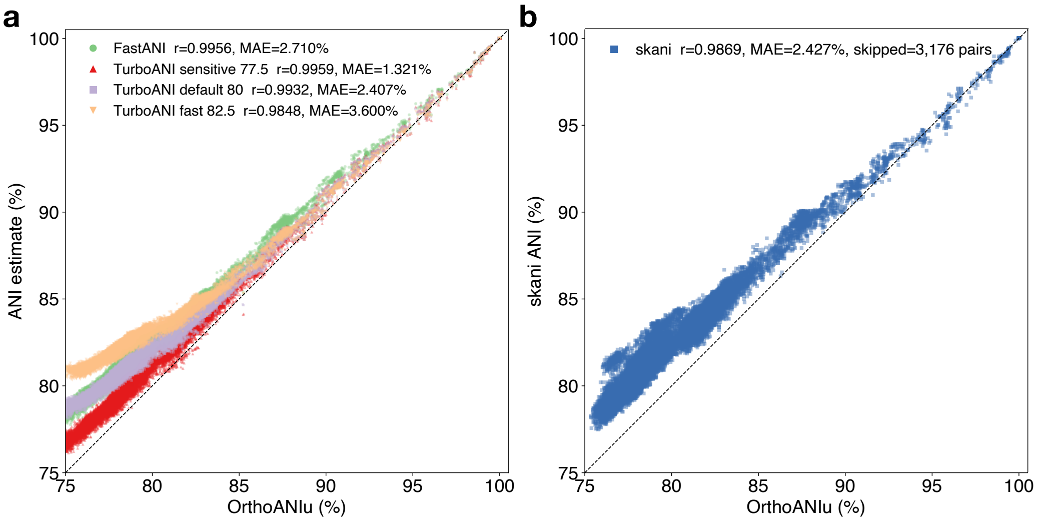


**Figure S5**. MAE and correlation between TurboANI 3 modes and orthoANIu (a), FastANI and orthoANIu (a) and skani and OrthoANIu (b).

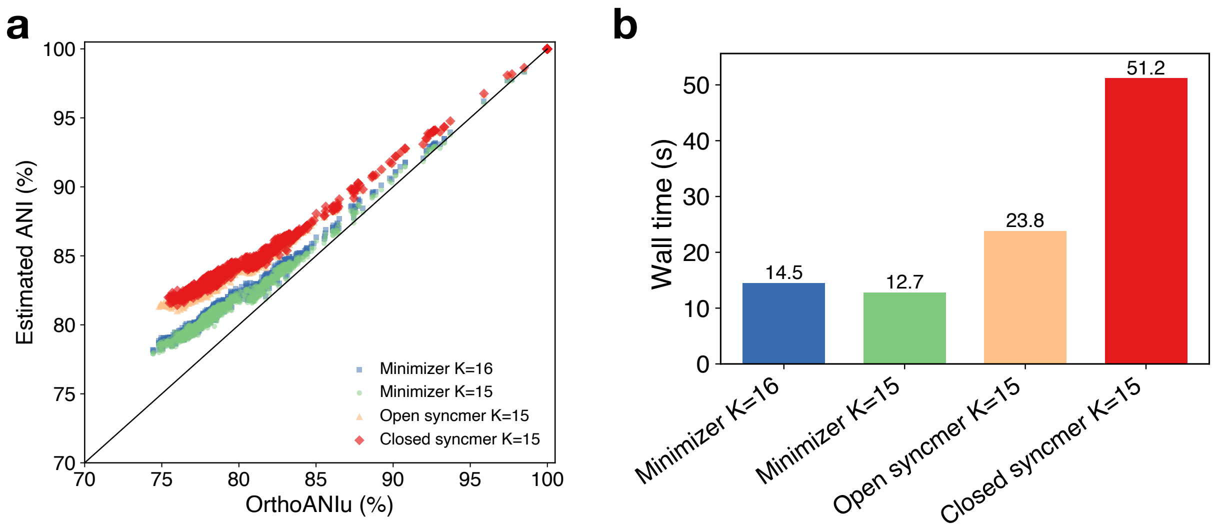


**Figure S6**. Open and closed syncmer-based TurboANI versus OrthoANIu truth (a) and running time comparisons with the default random minimizers (b).

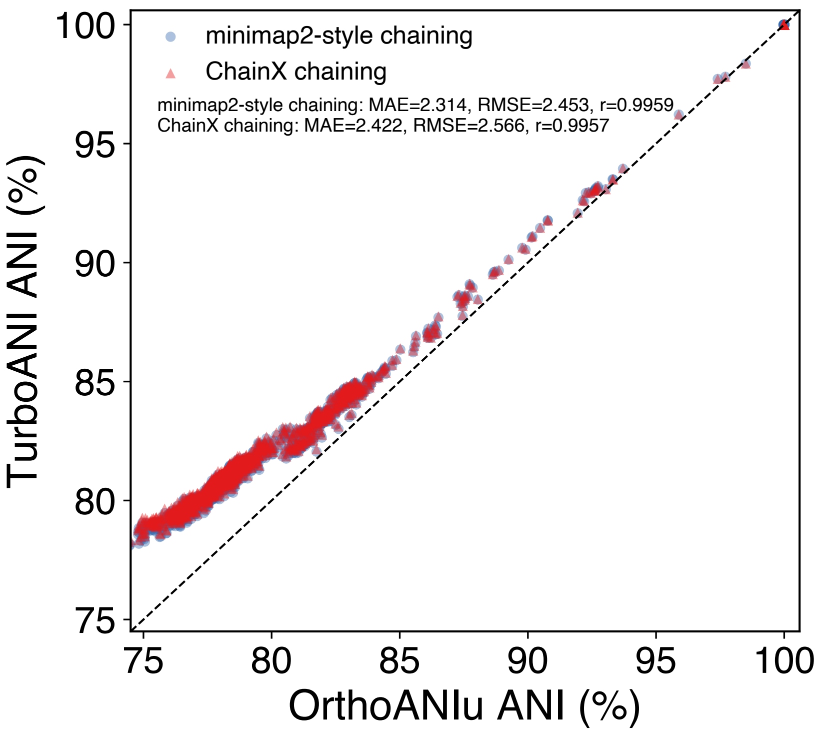


**Figure S7**. Optimal ChainX-based TurboANI estimation versus orthoANIu truth.

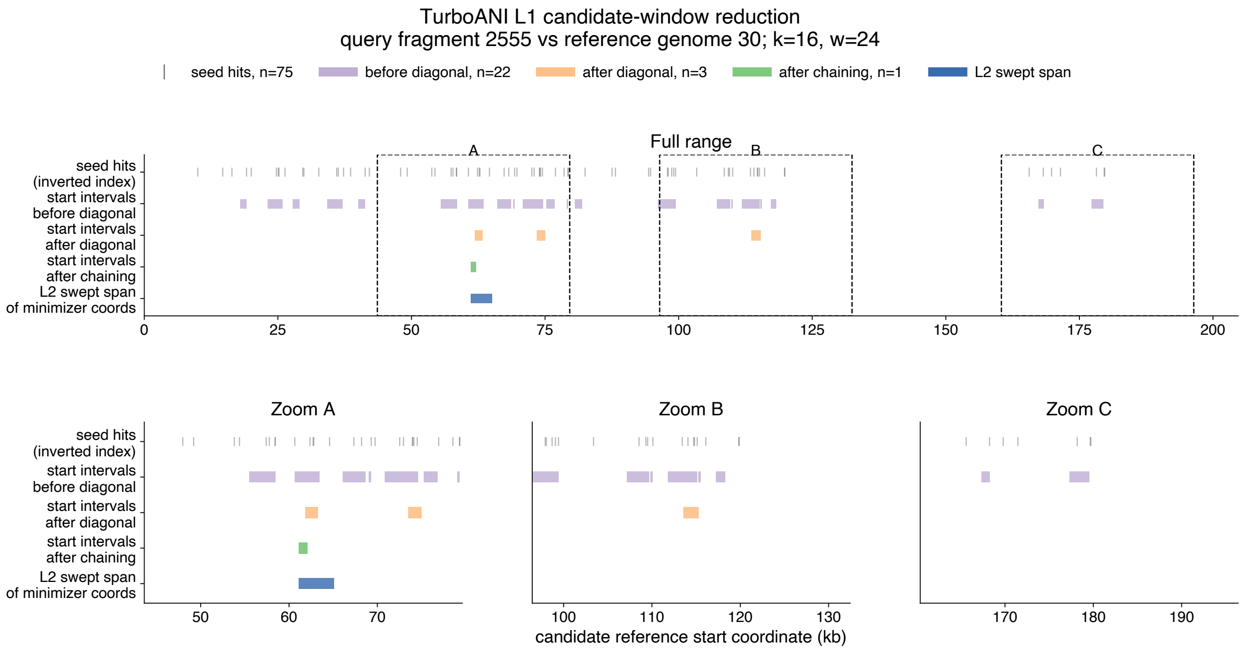


**Figure S8**. Detailed candidate interval changes for a query fragment during the L1 candidate interval screening process.

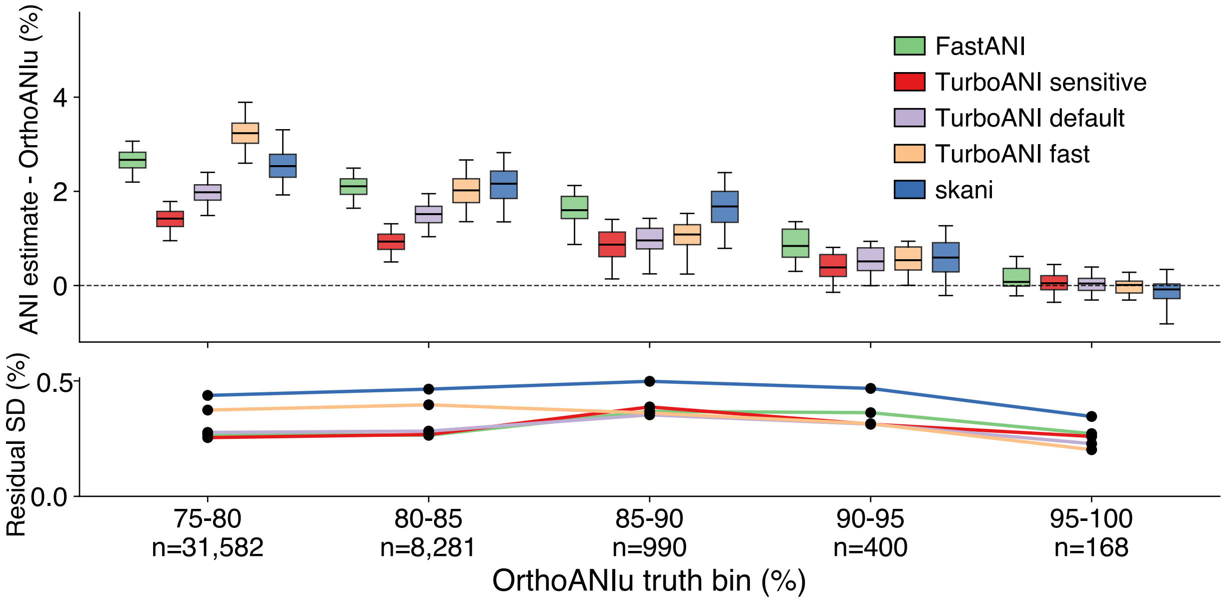


**Figure S9**. Mean error (showed as absolute ANI estimate – OrthoANIu truth) (a) and residual standard deviation (b) of each ANI estimation method (TurboANI 3 modes, FastANI and skani-sensitive) at 5 ANI bins from 75% to 100% ANI.

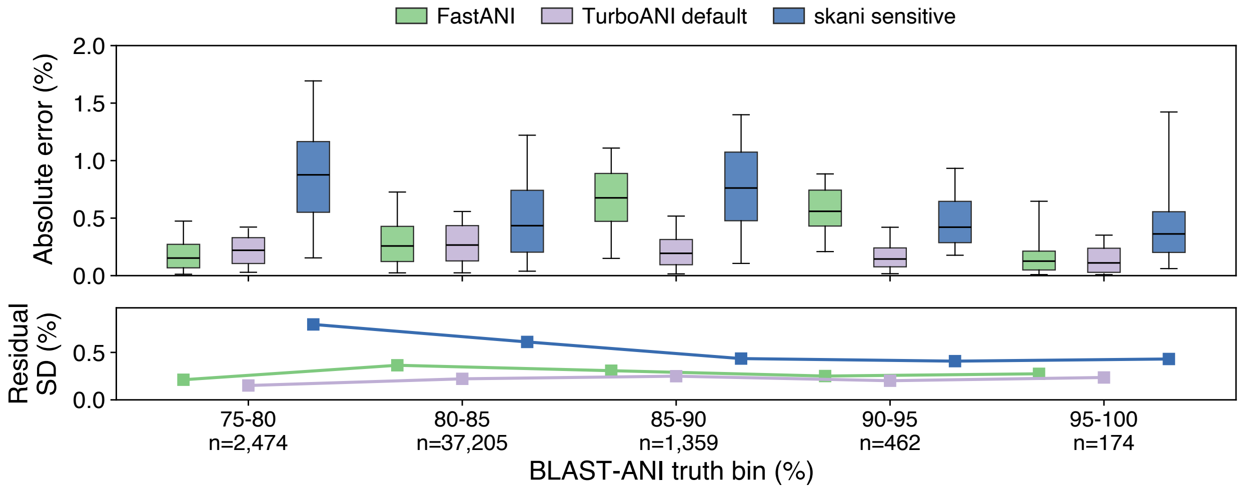


**Figure S10**. Mean error (showed as absolute ANI estimate – BLAST-ANI truth) (a) and residual standard deviation (b) of each ANI estimation method (TurboANI default mode, FastANI and skani-sensitive) at 5 ANI bins from 75% to 100% ANI.

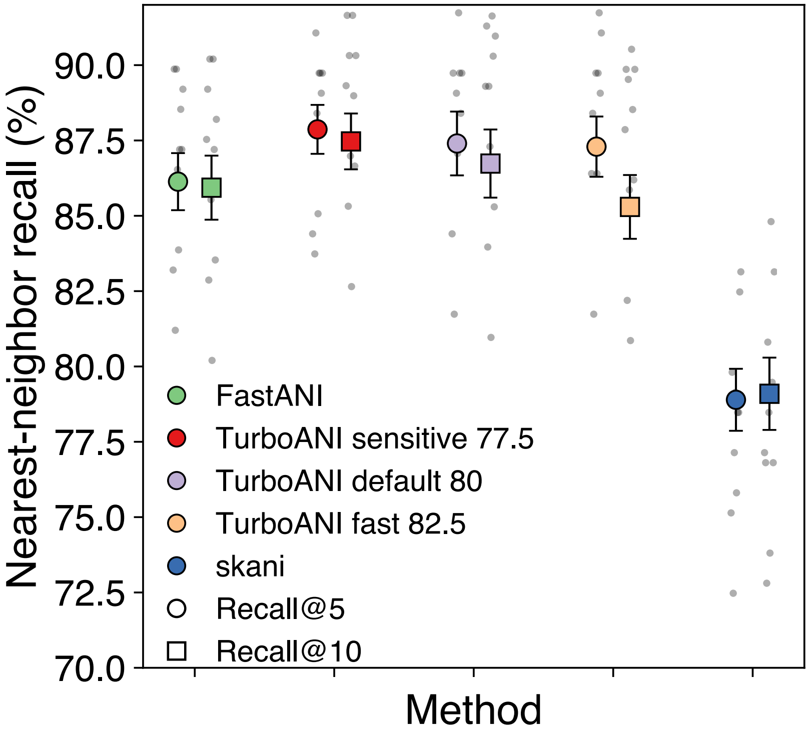


**Figure S11**. Nearest neighbor search accuracy benchmark for genome search and classification based on OrthoANIu. Top 10 and top 5 recalls were plotted for each ANI estimation method.

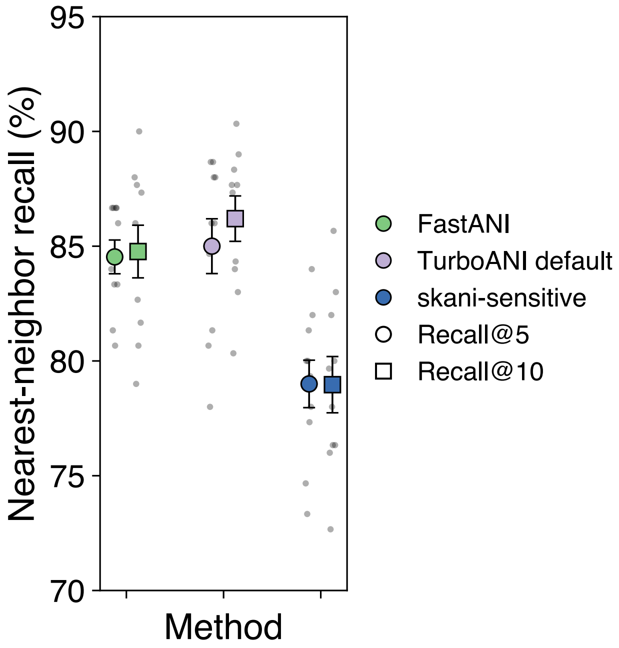


**Figure S12**. Nearest neighbor search accuracy benchmark for genome search and classification based on BLAST-ANI. Top 10 and top 5 recalls were plotted for each ANI estimation method.

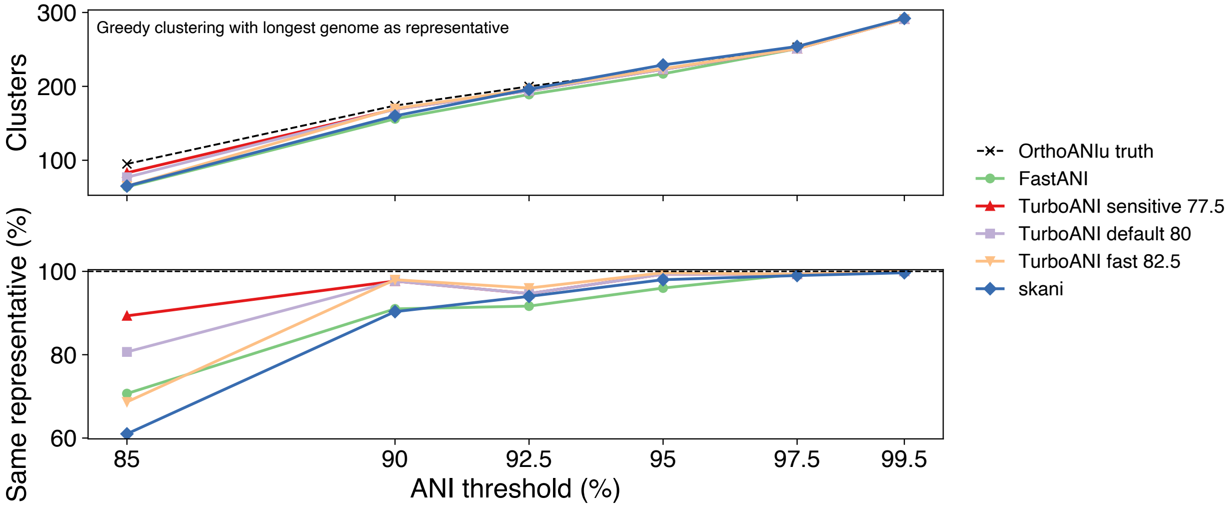


**Figure S13**. Greedy clustering accuracy benchmark for defining biological unit based on OrthoANIu. **a**. number of clusters at each clustering ANI threshold for each method. **b**. Representative recovery rate at each ANI clustering threshold for each method.

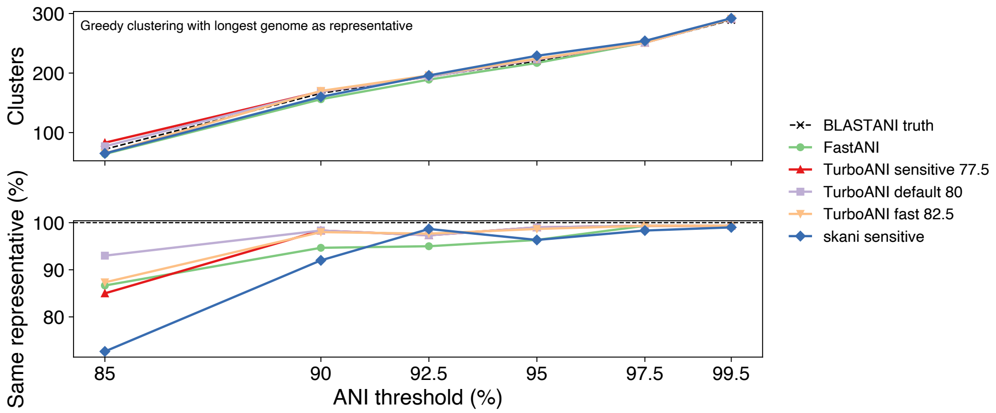


**Figure S14**. Greedy clustering accuracy benchmark for defining biological unit based on BLAST-ANI. **a**. number of clusters at each clustering ANI threshold for each method. **b**. Representative recovery rate at each ANI clustering threshold for each method.

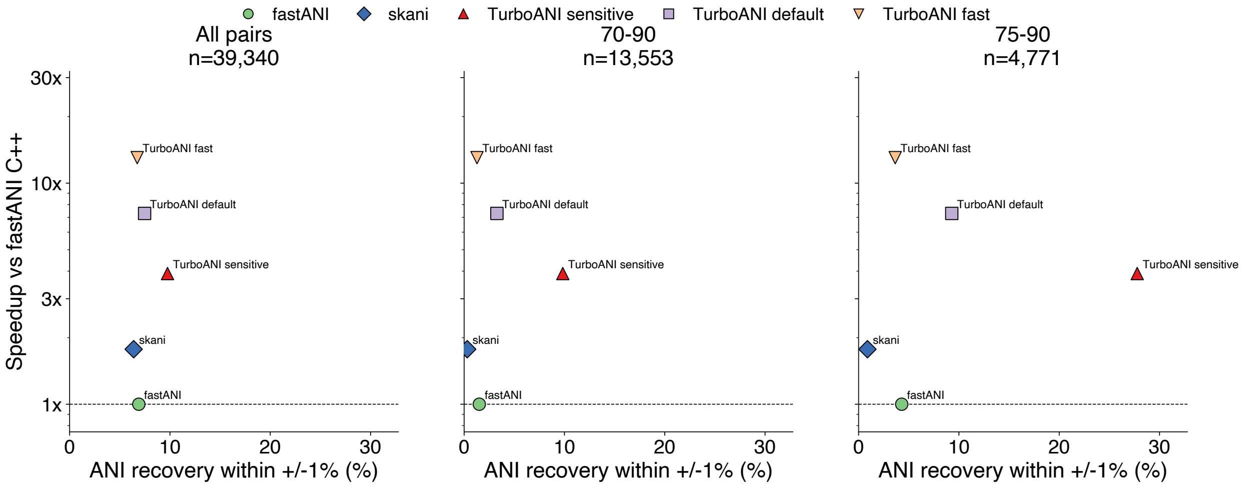


**Figure S15**. TurboANI, skani and FastANI speedup versus ANI recovery rate (within +/- 1% of the truth) against truth (OrthoANIu) for the archaeal family *Methanobacteriaceae*. (a) All ANI ranges, 75%~100%; (b) ANI range 75%~90%; (c) ANI range 75%~85%.

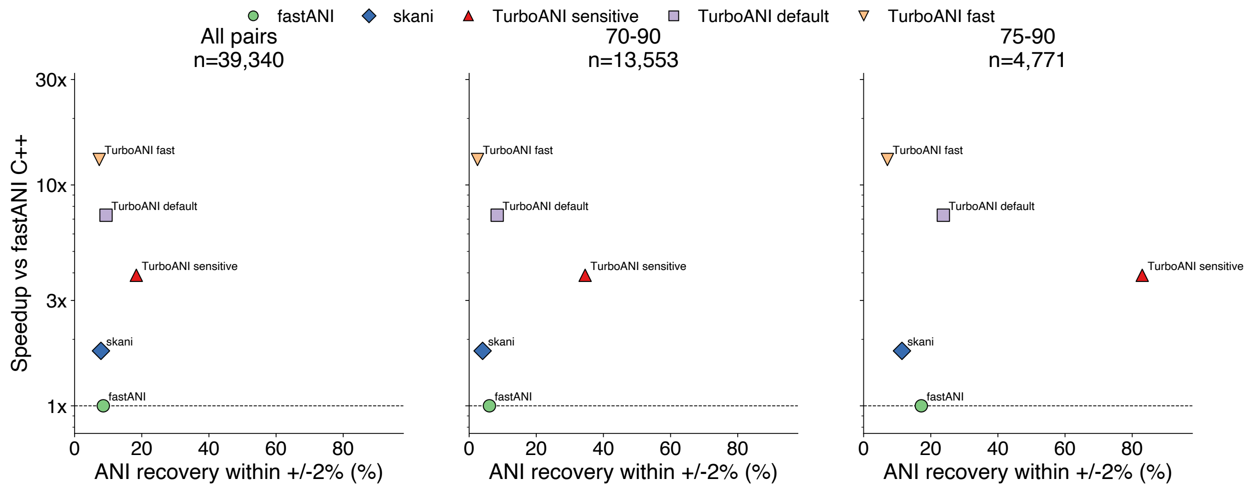


**Figure S16**. TurboANI, skani and FastANI speedup versus ANI recovery rate (within +/- 2% of the truth) against truth (OrthoANIu) for the archaeal family *Methanobacteriaceae*. (a) All ANI ranges, 75%~100%; (b) ANI range 75%~90%; (c) ANI range 75%~85%.

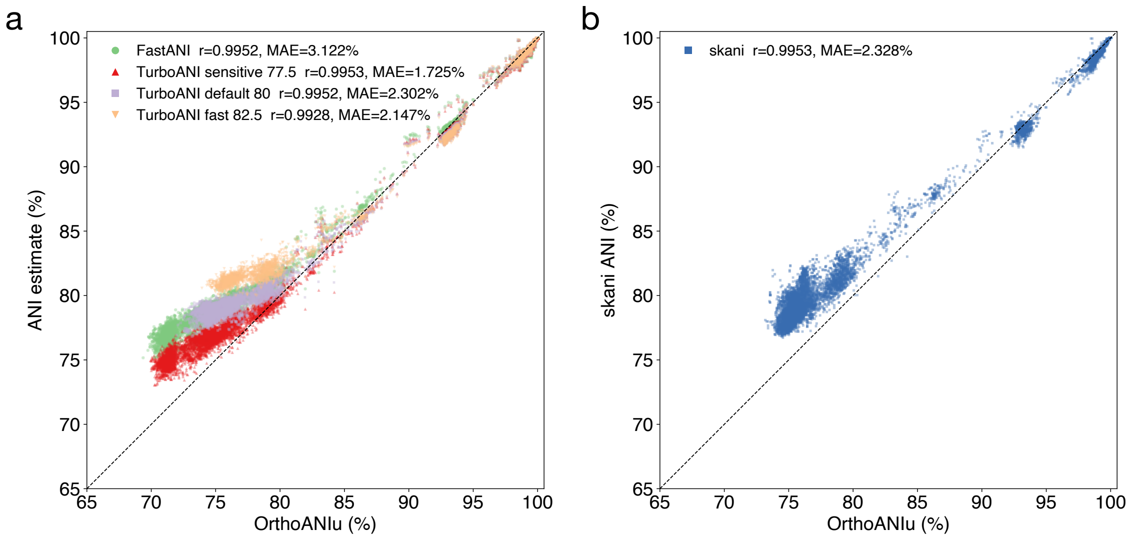


**Figure S17**. MAE and correlation between TurboANI 3 modes and orthoANIu (a), FastANI and orthoANIu (a) and skani and OrthoANIu (b). All comparisons are based on genomes from the archaeal family *Methanobacteriaceae*.

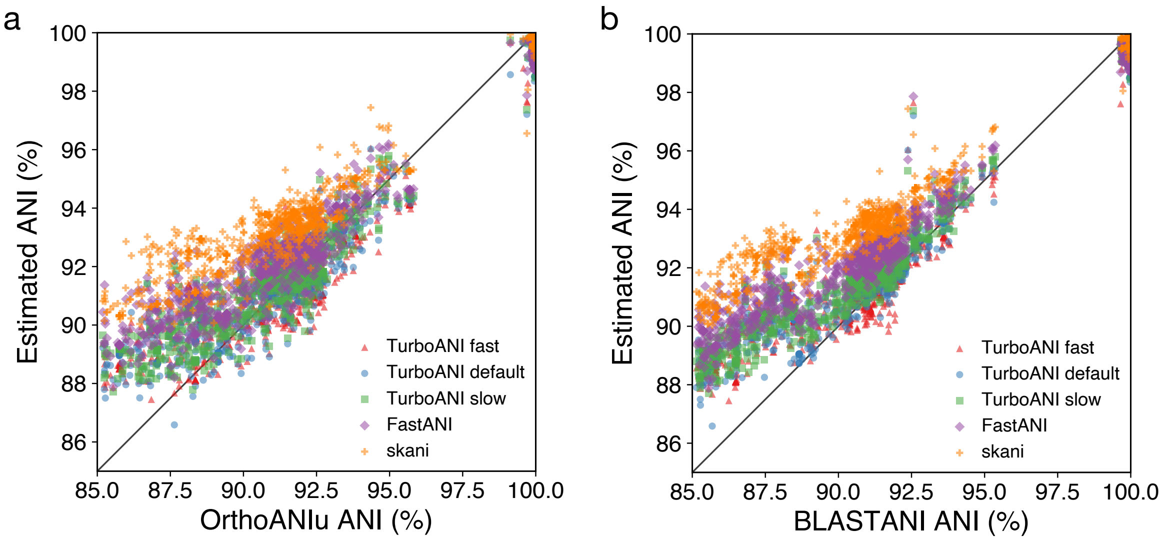


**Figure S18**. Correlation between TurboANI 3 modes, skani, FastANI and OrthoANIu (a) or BLAST-ANI (b) for virus genomes from the genus *Teseptimavirus*.

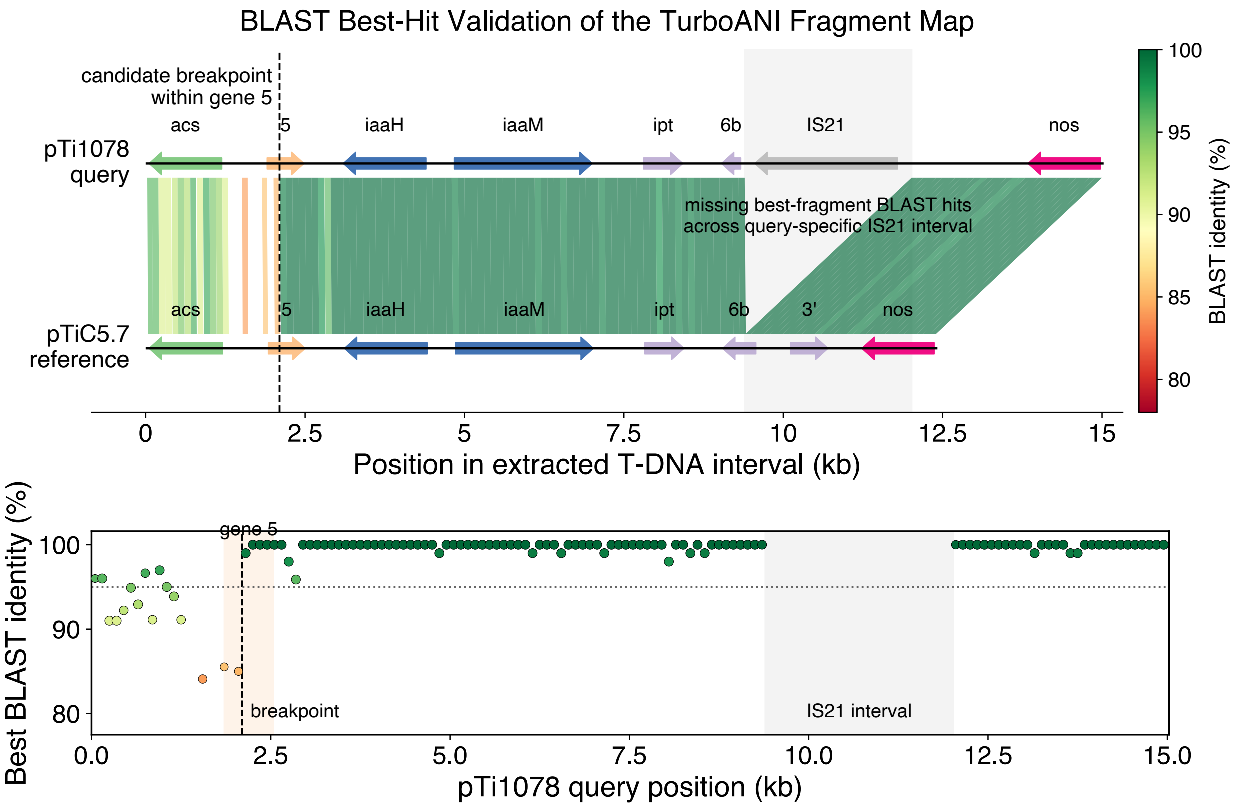


**Figure S19**. BLAST visualization of homologous mapping positions of the query Ti plasmid pTi1078 against the reference plasmid pTiC5.7. The best BLAST query matches were selected by bit scores after filtering by query alignment ratio (75%).

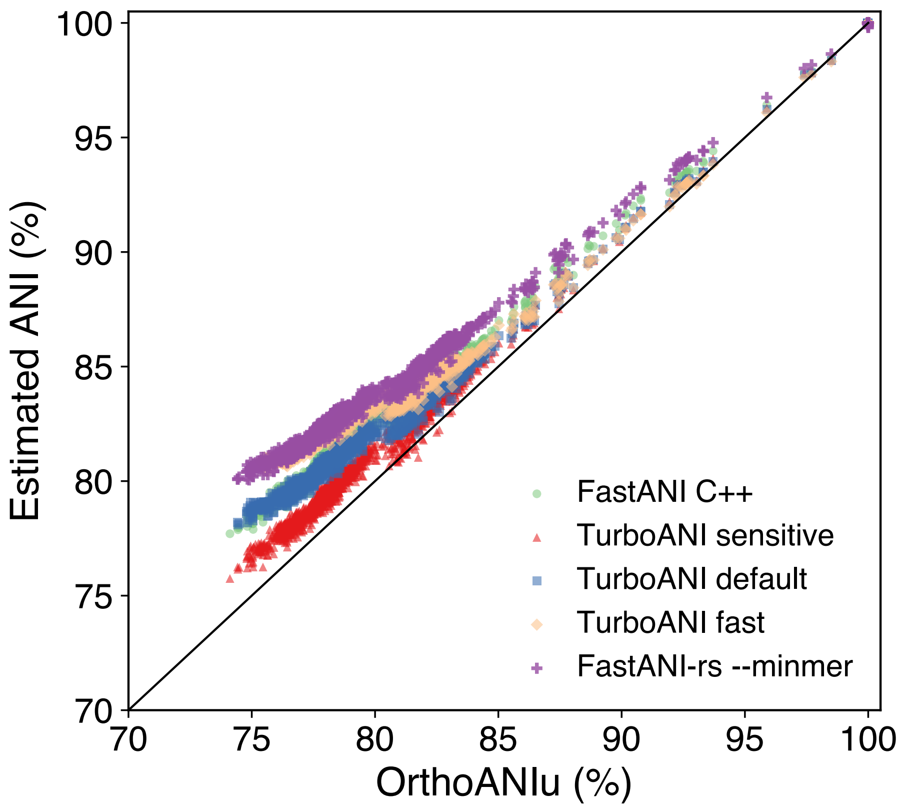


**Figure S20**. Minmer Jaccard estimator-based FastANI accuracy benchmark with original FastANI and TurboANI all modes. All ANI estimations were plotted against the OrthoANIu truth.

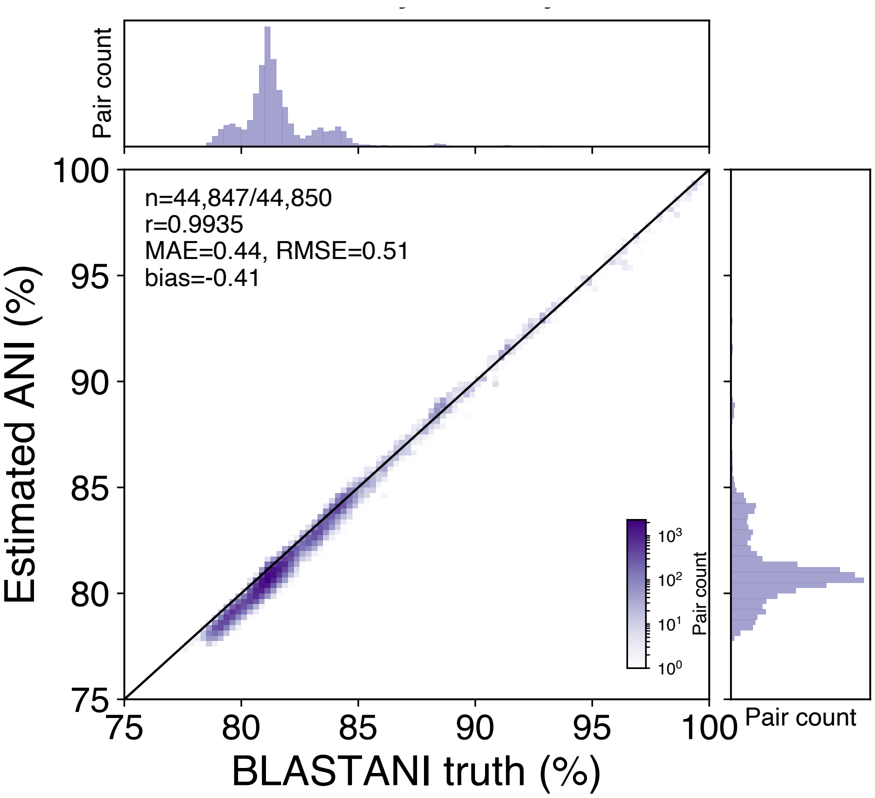


**Figure S21**. Correlation between TurboANI (without diagonal clustering and minimap2-style chaining) estimated ANI and BLAST-ANI truth for the 300-genome collection.

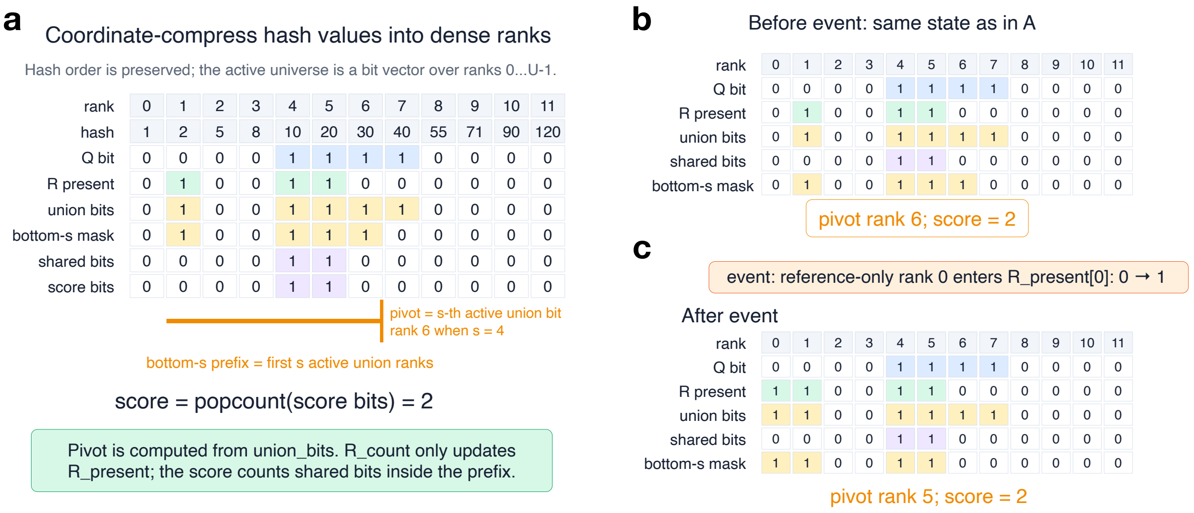


**Figure S22.** Bitset sliding MinHash algorithm. **a**. All query and reference minimizer hashes (the entire L2 interval) in the local L2 candidate interval are first sorted and replaced by dense ranks. Q marks ranks present in the query fragment, R_present marks rank whose reference count is currently nonzero in the sliding reference window, union_bits = Q OR R_present marks the ordered hash universe used for bottom-s selection, and shared_bits = Q AND R_present marks hashes shared by the query and the current reference window. The bottom-s sketch is represented by a prefix ending at the pivot rank, where the pivot is the rank of the s-th set bit in union_bits; therefore, all shared hashes contributing to the L2 score lie in shared_bits & prefix_mask. **b**. a simplified **a** without actual hash universe. **c**. When a reference hash enters or leaves the window, TurboANI updates only the affected reference count and flips the corresponding bit in R_present if the count changes between zero and nonzero. This may also flip the corresponding bit in union_bits and shared_bits. If the number of active union bits before the pivot changes, the pivot is repaired by walking to the previous or next set bit of union_bits until the prefix again contains exactly s active union ranks. The implementation finds the next set bit within a machine word using trailing_zeros() after masking off already-scanned lower bits, and finds the previous set bit using leading_zeros() function after masking off higher bits; a summary bit vector marks nonempty 64-bit blocks so that empty blocks can be skipped quickly. Once the pivot is repaired, the current L2 score is computed by a hardware popcount (count_one) over shared_bits & prefix_mask.

#### Supplementary Methods

**Detailed diagonal clustering followed by optimal ChainX**. For each query fragment, TurboANI first finds minimizer seed hits against the reference index, computes the seed diagonal $d = r - q$ and assigns each hit to a diagonal bin of width 1000 bp (default). Hits are grouped by (reference contig, diagonal bin), and only groups with at least $H_{min}$ unique query minimizer positions are considered. Here $H_{min}$ is derived from the relaxed FastANI minimum-hit criterion using the query fragment sketch size, k-mer length, minimum identity threshold, and distance model. For each retained diagonal group, each minimizer hit is treated as an anchor $a_i = (r_i, q_i, \ell)$, where $r_i$ is the reference minimizer position, $q_i$ is the query minimizer position, and $\ell = k$ is the k-mer length. TurboANI runs the chaining procedure twice, once in the forward query orientation and once in the reverse query orientation. In the forward orientation $q_i' = q_i$. In the reverse orientation, query coordinates are transformed as $q_i' = L_q - q_i - k$, where $L_q$ is the query fragment length. This allows decreasing query-coordinate seed hits to be represented as an increasing collinear chain.

The optimal ChainX dynamic program is implemented as a semi-global chaining problem with two sentinel anchors: a source anchor at and a sink anchor at $(-1, -1)$ and a sink anchor at $(L_r, L_q)$, where $L_r$ is the reference contig length and $L_q$ is the query fragment length. Anchors are sorted by (reference start, query start, length, sentinel status). The objective is to find a low-cost collinear chain from source to sink, with reference flanks effectively free but query overhangs penalized, so the chain is encouraged to explain the query fragment rather than merely a short local seed cluster. For two anchors $a_i$ and $a_j$, a transition is allowed only when $a_j$ strongly precedes $a_i$ in both reference and query coordinates: $r_j < r_i, r_j^{end} < r_i^{end}, q_j < q_i, q_j^{end} < q_i^{end}$. The transition cost is based on the larger unchained gap in reference or query, plus a correction for unequal overlap: $g_r = max(0, r_i - r_j^{end} - 1)$, $g_q = max(0, q_i - q_j^{end} - 1), o_r = max(0, r_j^{end} - r_i + 1), o_q = max(0, q_j^{end} - q_i + 1), c(j, i) = max(g_r, g_q) + |o_r - o_q|$ . For the sink transition, the reference terminal gap is set to zero, making the procedure semiglobal with respect to the reference. The implementation uses a sweep-line over anchor start and end events in reference-coordinate order. For each anchor start event, TurboANI considers predecessor anchors from three sources: the currently active anchor on the same diagonal, previous end events within the current distance bound, and active anchors on nearby diagonals whose diagonal distance is within the bound. A transition is accepted if it gives a lower cumulative cost; ties are resolved by preferring the longer chain. The initial distance bound is $B_0 = max(100, \lfloor 1.1(L_q - C_q) \rfloor)$, where $C_q$ is the total query-coordinate coverage of anchors in the diagonal group. If no feasible source-to-sink chain is found within the bound, the bound is multiplied by 4 and the dynamic program is rerun, up to 32 revisions. This follows the expanding-bound idea used by optimal ChainX chaining while keeping the practical implementation simple for TurboANI's relatively small per-bin anchor sets. TurboANI does not use only one best source-to-sink chain for L1 output. Instead, after the DP, it collects anchors that belong to any predecessor chain with at least $H_{min}$ chained anchors. This conservative union avoids discarding alternative plausible windows from the same diagonal group. The supported anchors are then converted into candidate reference-window starts using $s_i = r_i - q_i'$, deduplicated, expanded by diagonal band (500 bp) on both sides, and merged if adjacent or overlapping on the same reference contig.
